# Surfaceome CRISPR Screen Identifies OLFML3 as a Rhinovirus-inducible IFN Antagonist

**DOI:** 10.1101/2020.11.08.372607

**Authors:** Hong Mei, Zhao Zha, Wei Wang, Yusang Xie, Yuege Huang, Wengping Li, Dong Wei, Xinxin Zhang, Jia Xie, Jieming Qu, Jia Liu

## Abstract

**Background:** Rhinoviruses (RVs) cause more than half of common cold and, in some cases, more severe diseases. Functional genomics analyses of RVs using siRNA or genome-wide CRISPR screen uncovered a limited set of host factors, few of which has proven clinical relevance.

**Results:** Herein, we systematically compared genome-wide CRISPR screen and surface protein-focused CRISPR screen, referred to as surfaceome CRISPR screen, for their efficiencies in identifying RV host factors. It was found that surfaceome screen outperformed genome-wide screen in the success rate of hit identification. Importantly, using surfaceome screen we have identified olfactomedin like 3 (OLFML3) as a novel host factor of RV serotypes A and B including a clinical isolate. We found that OLFML3 was a RV-inducible suppressor of the innate immune response and that OLFML3 antagonized type I interferon (IFN) signaling in a SOCS3-dependent manner.

**Conclusion:** Our study has suggested that RV-induced OLFML3expression is an important mechanism for RV to hijack the immune system and underscored surfaceome CRISPR screen in identifying viral host factors.

## Background

The emerging genome engineering technologies, including clustered regularly interspaced short palindromic repeats (CRISPR)-CRISPR-associated protein 9 (Cas9), have transformed basic and translational biomedical research [1]. In particular, functional genomics using CRISPR screen provides unprecedented approaches to establishing and understanding phenotype-genotype relationships [2]. For example, CRISPR screen has been widely used to identify and dissect the cellular host factors for a variety of viruses[3] including noroviruses [4], human immunodeficiency virus (HIV) [5], flaviviruses [6, 7], influenza viruses [8, 9], picornaviruses [10–12], alphaviruses [13, 14] and others.

RVs are known as the prevalent pathogen causing common cold [15] and have also been found to be associated with other severe respiratory symptoms including asthma exacerbations [16] and chronic obstructive pulmonary disease [17]. Despite the increasing number of RV-associated severe respiratory diseases, the causal link between RV infection and clinical outcome remains poorly understood. Particularly, the diverse categories of RVs make it extremely sophisticated to dissect host-pathogen interactions. Functional genomics has been employed to understand RV infections, including RNAi or haploid cell-based genetic screen [18, 19] and emerging CRISPR screens [3]. However, conventional genome-wide genetic screen appeared to have limited efficiency and only a few novel host factors of RVs have been identified and validated to have clinical relevance [12, 18].

It has been shown that cell proliferation and cell cycle-related proteins such as p53 can perturb the outcome of genome-wide CRISPR screen by masking the positive hits [20]. One strategy to overcome this problem is to employ focused screening. To date, a number of focused CRISPR libraries have been constructed to realize genetic screen on kinome [21, 22], epigenome [23, 24] and cancer-related [25, 26] genes. Importantly, cell surface protein-focused CRISPR libraries have been explored in previous studies [11, 27, 28]. However, these studies did not systematically analyze the efficiencies of genome-wide CRISPR screen and surfaceome CRISPR screen. A thorough comparative study can elucidate the difference between genome-wide and surfaceome screens for their efficiencies in identifying viral host factors.

In the present study, we constructed genome-wide and surfaceome CRISPR libraries using identical algorithms and performed genome-wide and surfaceome CRISPR screens in parallel to screen for RV host factors. In contrast to the low success rate of genome-wide CRISPR screen, surfaceome screen identified a set of cell surface host factors that were important for RV infection. Notably, OLFML3 was found to be an RV-inducible dependency factor that promoted the infection. We showed that OLFML3 suppressed the innate immune response of host cells via SOCS-mediated negative regulatory pathway of type I IFN signaling.

## Results

### Design of sgRNA for genome-wide and surfaceome CRISPR-Cas9 libraries

We constructed CRISPR libraries using previously established algorithms for evaluating on-target [29] and off-target [30] activities. The genome-wide library contained 18,421 genes with 12 sgRNA for each gene and was divided into three sub-libraries A, B and C according to the scores of sgRNA (Additional file 1: Fig. S1a and Additional file 3). The collection of cell surface proteins was defined using a mass spectrometry (MS) database that investigated cell surface proteins in 41 individual human cells [31]. The union set contains 1,344 genes and the 12 sgRNA targeting to these genes were extracted from the genome-wide library to construct the surfaceome library (Additional file 1: Fig. S1a and Additional file 4). Cross reference of the 1,344 cell surface proteins by gene ontology dataset for cellular component showed that the majority of these proteins are located on plasma membrane and/or extracellular matrix (Fig. 1a).

**Fig. 1.**
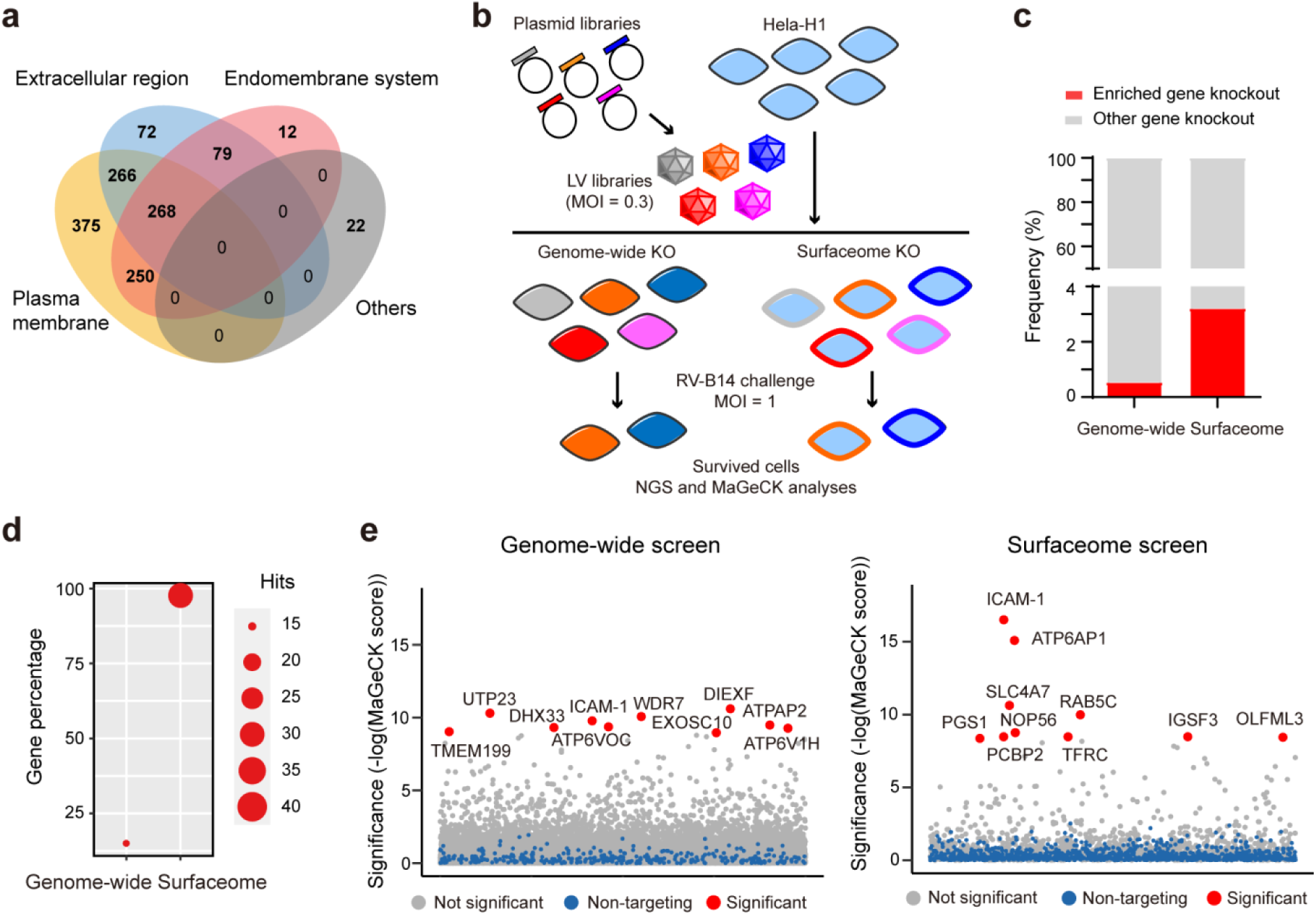
Genome-wide and surfaceome-wide screens for identification of host factors of RV. **a** Venn diagram showing the composition of surfaceome library. **b** Flow chart showing the procedures of genome-wide and surfaceome CRISPR screens. **c** Analyses of enriched sgRNA, or gene knockout, in genome-wide and surfaceome screens. **d** Analyses of the compositions of positive hits, as defined by a FDR of less than 0.01, in genome-wide and surfaceome screens. **e** Bubble plot showing the results of CRISPR screen. Top 10 candidate hits are shown. Significance of enrichment was calculated by MAGeCK.

### Construction of genome-wide and surfaceome CRISPR-Cas9 libraries in H1-Hela cells

To normalize for the quality of sgRNAs in the library, we performed side-by-side screening with RVs using genome-wide sub-library A and surfaceome library, which exhibited similar on-target and off-target scores (Additional file 1: Fig. S1b). This sub-library A contains 73,527 sgRNA targeting to 18,421 genes with an average of four sgRNA per gene. In comparison, surfaceome CRISPR screen involved 16,975 sgRNA targeting to 1,344 genes. These libraries were packaged into lentiviruses (LVs) and transduced with a multiplicity of infection (MOI) of 0.3 into H1-Hela cells that are known to support RV infection [32] (Fig. 1b). Next-generation sequencing (NGS) analyses of the PCR amplicons of genome-integrated sgRNA revealed that plasmid and cell libraries of genome-wide and surfaceome screens had full coverage of sgRNA with optimum distribution, as evidenced by Gini coefficients of less than 0.1 [33] (Additional file 1: Fig. S2a). The cumulative frequency of sgRNA reads in these four libraries were consistent with the predicted distribution (Additional file 1: Fig. S2b-c). Collectively, these results validated the quality of the constructed CRISPR libraries for subsequent RV challenge.

### Identification of RV host factors on H1-Hela using genome-wide and surfaceome CRISPR-Cas9 libraries

To identify RV host factors, we performed the screen using a previously reported RV-induced cell death model [32]. H1-Hela cells harboring genome-wide or surfaceome CRISPR library were challenged with RV serotype B14 (RV-B14) at an MOI of 1.0 for 48 h. Upon completion of RV-B14 challenge, survived cells were collected and sgRNAs were PCR-amplified and then analyzed by NGS. MAGeCK analyses [33] of the RV infection groups in comparison with mock groups (Fig. 1b) identified a set of enriched sgRNA (Additional file 1: Fig. S3a). Using a false discovery rate (FDR) of less than 0.01 as a cutoff for sgRNA enrichment, surfaceome screen identified 3.2% of the total 1,344 surface proteins bearing enriched sgRNA, while genome-wide screen uncovered sgRNA enrichment in 0.54% of the 18,421 genes in the library (Fig. 1c). Of the enriched sgRNA in the genome-wide screen, less than 20% were targeted to the surface proteins (Fig. 1d). It was noted that surfaceome screen identified more absolute number of enriched sgRNA for surface proteins than genome-wide screen (Fig. 1d).

Next we ordered the genes with enriched sgRNA using robust rank aggregation (RRA) analyses in the MAGeCK pipeline (MAGeCK score) and displayed top 10 candidate hits in each library (Fig. 1e and Additional files 5-6). Both genome-wide and surfaceome screens identified intercellular adhesion molecule 1 (ICAM-1), a known receptor for RV serotypes A and B[34], in the top 10 hits (Fig. 1e). Except for ICAM-1, these screens resulted in no overlap in the top 10 hits. It has to be noted that the surfaceome and genome-wide CRISPR libraries contain 12 and 4 sgRNA respectively for each gene. To exclude the possibility that the differential hits in genome-wide and surfaceome screens were due to the effects of different number of sgRNAs, we performed a pseudo screen with the surfaceome library where sgRNA 1 to 4 in the sub-library A (Additional file 1: Fig. S1), rather than all the 12 sgRNA, were included for each surface protein. MAGeCK analyses of the pseudo screen revealed a generally consistent rank of the top 10 hits with the experimental screen (Additional file 1: Fig. S3b). Collectively, our results suggested that both genome-wide and surfaceome screens were robust enough to identify strong candidates such as RV receptor and that surfaceome screen was more efficient in identifying candidate hits for surface proteins.

### Validation of the top 10 candidate hits in genome-wide and surfaceome CRISPR screens

Next we sought to validate the top 10 hits from genome-wide and surfaceome screens by constructing knockout cell lines for individual genes. Two sgRNA were designed for each gene and the gene disruption efficiency was evaluated by T7E1 assay (Additional file 1: Fig. S4). Non-targeting sgRNA was used as mock for subsequent phenotypical analyses. In similar experimental settings with the virus challenge screen, we examined RV-induced cell death on individual cell lines of the top 10 hits identified from surfaceome and genome-wide screens. It was found that in the genome-wide screen only ICAM-1 knockout exhibited consistently improved cell viability with both sgRNAs compared to non-targeting sgRNA, whereas surfaceome screen identified 6 gene knockout out that displayed protective effects against RV-14 infection (Fig. 2a). Next we analyzed the effects of knockout of the identified genes on viral replication. Individual knockout cells were infected with RV-14 at an MOI of 2 and viral RNA was extracted from medium supernatant or cell lysates at 24 h post infection. Among the top 10 hits from genome-wide screen, only ICAM-1 knockout resulted in significantly reduced viral loads in medium supernatant in comparison with non-targeting sgRNA (Fig. 2b). Investigation of viral loads in cell lysates uncovered generally consistent results (Additional file 1: Fig. S5). These results suggested that except for ICAM-1 the top 10 candidate genes identified by genome-wide screen were most likely false positive hits. By contrast, the 6 gene knockout from surfaceome screen that exhibited protective effects (Fig. 2a) showed consistently reduced viral loads in medium supernatant (Fig. 2b). The function of these 6 genes was further examined by immunofluorescence staining of RV-B14 envelope protein for evaluation of the effects of gene knockout on viral infection rate. It was found that in addition to ICAM-1, the knockout of IGSF3, RAB5C and OLFML3 showed consistently reduced RV-B14 infection rates with both sgRNA at an MOI of 1 or 2 (Fig. 2c and Additional file 1: Fig. S6). Collectively, these results indicated that surfaceome CRISPR screen was more efficient than genome-wide screen in identifying RV host factors.

**Fig. 2.**
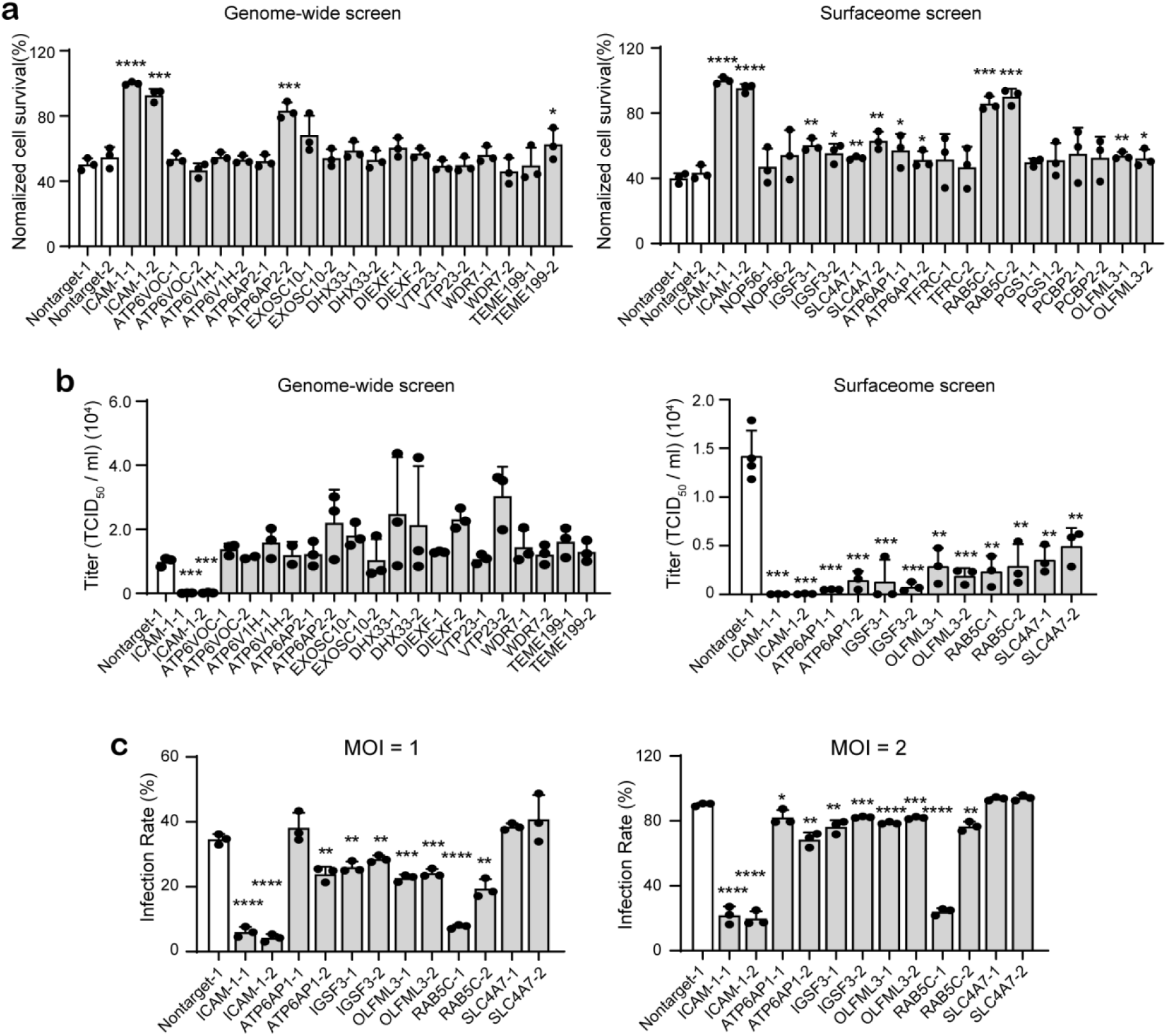
Validation of the top 10 hits from surfaceome and genome-wide screens. **a** Cell viability assay for determination of the protective effects of identified gene knockout on RV-B14-induced cell death. The assay is performed at 24 h post infection of RV-B14 at an MOI of 2. **b** RT-qPCR quantification of viral loads in medium supernatant. The supernatant is harvested at 24 h post infection of RV-B14 at an MOI of 2. **c** Immunofluorescence (IF) staining of RV-B14 envelop protein for evaluation of infection rates at individual cell lines. IF staining is performed at 16 h post infection of RV-B14 at an MOI of 1 or 2. Each biological replicate contains the quantification results from 2,000 cells. The significant difference between knockout cells and non-targeting sgRNA mock groups are determined using two-tailed unpaired Student’s *t*-test. *, *P* <0.05; **, *P* <0.01; ***, *P* <0.001; ****, *P* <0.0001.

### Validation of RAB5C and OLFML3 as RV dependency factors

Although a series of candidate dependency factors were identified for RV-B14 from surfaceome screen, we chose to focus subsequent analyses on RAB5C and OLFML3. RAB5C was reported to involve in the entry process of flaviviruses [35]. OLFML3 was found to function as an immunosuppressive molecule during the carcinogenesis of glioblastoma [36]. These two proteins may represent novel dependency factors of RV-B14, none of which have been reported to involve in RV infection. As a known receptor for RV serotypes A and B [37, 38], ICAM-1 was included as positive control for the following experiments.

Single clones of ICAM-1, RAB5C and OLFML3 knockout cells were constructed and validated (Additional file 1: Fig. S7). It was found that ICAM-1^-/-^, RAB5C^-/-^ and OLFML3^-/-^ cells all exhibited significantly increased cell viability upon RV-B14 challenge compared with non-targeting sgRNA-treated H1-Hela cells (Additional file 1: Fig. S8a-b). These knockout cells showed markedly reduced viral loads in medium supernatant (Fig. 3a) and cell lysates (Fig. 3b) over monitored time course, where ICAM-1 or RAB5C knockout nearly abolished viral replication (Fig. 3a-b). To further validate the importance of RAB5C and OLFML3 for RV infection, we performed rescue experiments by overexpressing these genes in corresponding knockout cells. It was found that RAB5C and OLFML3 overexpression rescue could restore RV-B14-induced cell death in H1-Hela knockout cells (Fig. 3c). Importantly, consistent results were observed with a different RV serotype RV-A16 (Fig. 3d). RAB5C and OLFML3 overexpression also rescued the replication of RV-B14 or RV-A16 in corresponding knockout cells, as determined by viral loads in medium supernatant or cell lysates (Fig. 3e-h). Similarly, ICAM-1 overexpression could rescue the susceptibility of ICAM-1^-/-^ cells to RV infection (Additional file 1: Fig. S8c-f). Collectively, these results demonstrate that RAB5C and OLFML3 are host dependency factors of RV.

**Fig. 3.**
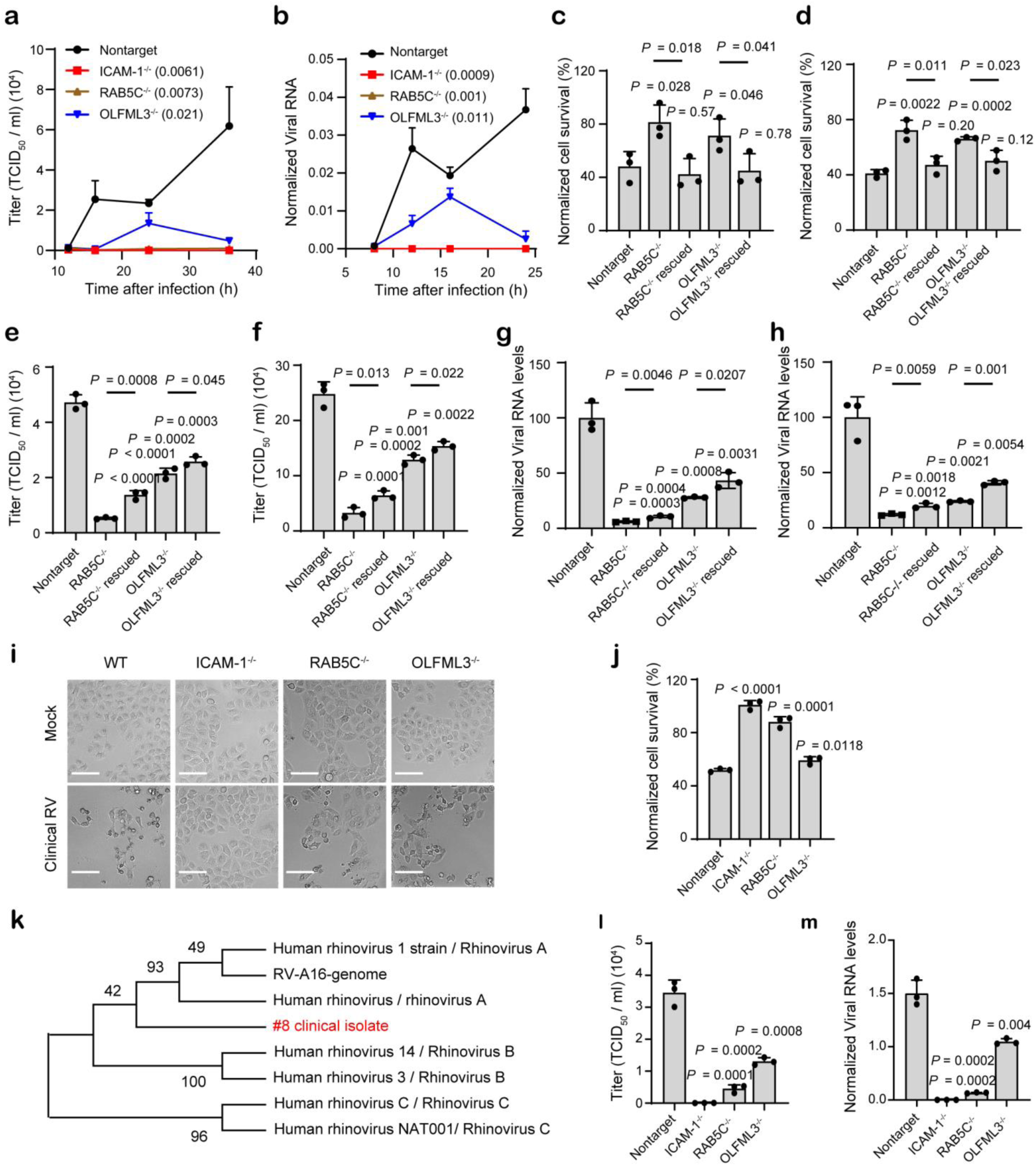
Validation of the effects of ICAM-1, RAB5C and OLFML3 on RV infection. **a-b** Time-dependent viral replication of RV-B14 in mock and knockout cells, as determined by viral loads in medium supernatant (**a**) and cell lysate (**b**). Cells are infected with RV-B14 at an MOI of 2. Viral RNA in cell lysates (**b**) is normalized to RPLP0 expression. Significant difference between test groups and non-targeting sgRNA group is determined using two-way ANOVA with Dunnett’s multiple comparisons test. **c**-**d** Rescued susceptibility of knockout cells to the infection of RV-B14 (**c**) and RV-A16 (**d**) by overexpression of RAB5C and OLFML3 respectively. **e-f** Rescued replication of RV-B14 (**e**) and RV-A16 (**f**) in knockout cells by overexpression of RAB5C and OLFML3 respectively, as determined by viral loads in medium supernatant. **g-h** Rescued replication of RV-B14 (**g**) and RV-A16 (**h**) in knockout cells by overexpression of RAB5C and OLFML3 respectively, as determined by viral loads in cell lysates. Viral RNA in cell lysates is normalized to RPLP0 expression. **i** Representative images of CPEs induced by clinical RV strain. Scale bar, 100 μm. **j** Cell viability of mock and knockout cells upon challenge of clinically isolated RV strain. **k** Phylogenetic analyses of clinical RV strain using MEGA X [50], with VP4 gene as the reference. **l**-**m** The effects of gene knockout on the replication of clinical RV strain, as determined by RT-qPCR quantification of viral loads in medium supernatant (**l**) or cell lysates (**m**). For **m**, Viral RNA in cell lysates is normalized to RPLP0 expression. For **c-j** and **l**-**m**, analyses are performed at 24 h post infection of RV at an MOI of 2. Significant difference between test groups and non-targeting sgRNA group is determined using two-tailed Student’s *t* test and the *P* values are shown. Significant difference between knockout and overexpression rescue groups is determined using two-tailed Student’s *t* test and the *P* values are shown above the lines.

Next we sought to examine the effects of RAB5C or OLFML3 knockout on the infection of clinical RV strain. A RV strain was isolated from the oropharyngeal swab of a patient with upper respiratory tract infection. This RV isolate induced notable cytopathic effects (CPEs) in H1-Hela cells (Fig. 3i) as laboratory strains did. ICAM-1^-/-^, RAB5C^-/-^ and OLFML3^-/-^ H1-Hela cells all exhibited significant resistance to the clinical RV strain (Fig. 3i-j). Phylogenetic analyses using the VP4 gene suggested that the clinical RV isolate was closely related with the serotype A RV with less similarity with RV-B and RV-C strains (Fig. 3k). Further investigation of viral loads in medium supernatant (Fig. 3l) and cell lysates (Fig. 3m) suggested that ICAM-1, RAB5C or OLFML3 knockout abolished or inhibited the replication of the clinical RV strain. These results thus verified the consistent functions of RAB5C and OLFML3 during the infection of laboratory and clinical RV strains.

### Dissection of the functions of RAB5C and OLFML3 in RV infection

To analyze the functions of RAB5C and OLFML3, we first sought to determine whether these proteins were involved in virus attachment and entry. For virus attachment assay, RV-B14 was incubated with mock and knockout cell lines at 4 ℃ and cells with attached virus were harvested and lysed for RT-qPCR quantification of viral loads. Consistent with the known function of ICAM-1 as the receptor of RV serotypes A and B, ICAM-1^-/-^ cells exhibited significantly reduced virus attachment. In contrast, knockout of RAB5C or OLFML3 did not lead to reduced virus attachment (Additional file 1: Fig. S9a). For virus entry assay, RV-B14 was incubated with mock and knockout cell lines first at 4 ℃ and then the internalization of RV-B14 was initiated by incubation at 37 ℃. Surface-bound RV-B14 was removed by extensive washing and internalized virus was quantified by RT-qPCR. It was found that RAB5C^-/-^ or OLFML3^-/-^ cells did not show reduced virus entry (Additional file 1: Fig. S9b). Consistent results were observed with the functions of RAB5C and OLFML3 in RV-A16 attachment and entry (Additional file 1: Fig. S9c-d). Fluorescence in situ hybridization (FISH) detection of internalized RV-B14 showed similar findings for the roles of RAB5C and OLFML3 in virus entry (Additional file 1: Fig. S9e-f). These results together suggested that RAB5C or OLFML3 did not affect RV attachment or entry.

Next we sought to analyze whether RAB5C and OLFML3 were involved in the life cycle of RV after uncoating of viral genome. The RNA encoding RV-A16 genome was transfected into cells to bypass virus entry and uncoating processes. It was found that mock and knockout cells exhibited similar cell viability (Additional file 1: Fig. S9g) and viral loads (Fig. 4a-b). This suggested that RAB5C or OLFML3 did not involve in the processes after genome uncoating. This experiment along with the above virus attachment and entry assays excluded the functions of RAB5C or OLFML3 in many processes during RV replication. In order to investigate whether RAB5C and OLFML3 participated in RV genome RNA uncoating, we used a previously established guanidine hydrochloride (GuHCl)-mediated mRNA synthesis suppression assay [12] to analyze the dynamics of intracellular viral RNA. Mock and knockout H1-Hela cells displayed comparable cell viability in the presence of 2 mM GuHCl (Additional file 1: Fig. S9h). During the monitored time course of 24 h, RV-B14 RNA increased in a time-dependent manner in wild-type H1-Hela cells. By contrast, treatment of cells with 2 mM GuHCl led to notable time-dependent RNA decay (Additional file 1: Fig. S9i), suggesting of inhibited synthesis of viral RNA and active cellular RNA degradation machinery. In the presence of 2 mM GuHCl, RV-B14 and RV-A16 infection in ICAM-1^-/-^ cells resulted in little viral loads in cell lysates at all time points (Fig. 4c-d), which was in consistency with the above results of ICAM-1^-/-^ cells (Fig. 3a-b). With 2 mM GuHCl treatment, the dynamics of viral RNA in OLFML3^-/-^ cells displayed similar pattern to that of non-targeting sgRNA-treated cells, while RAB5C^-/-^ cells had significantly higher viral RNA level at 3 and 6 h post RV infection (Fig. 4c-d). Given inhibited RNA synthesis, the slower decay rate of viral RNA in RAB5C^-/-^ cells could be attributed to the protection of viral RNA from cellular RNA degradation machinery as a result of perturbed uncoating process of viral genome. Collectively, these results suggested that RAB5C, but not OLFML3, may be involved in RV genome uncoating.

**Fig. 4.**
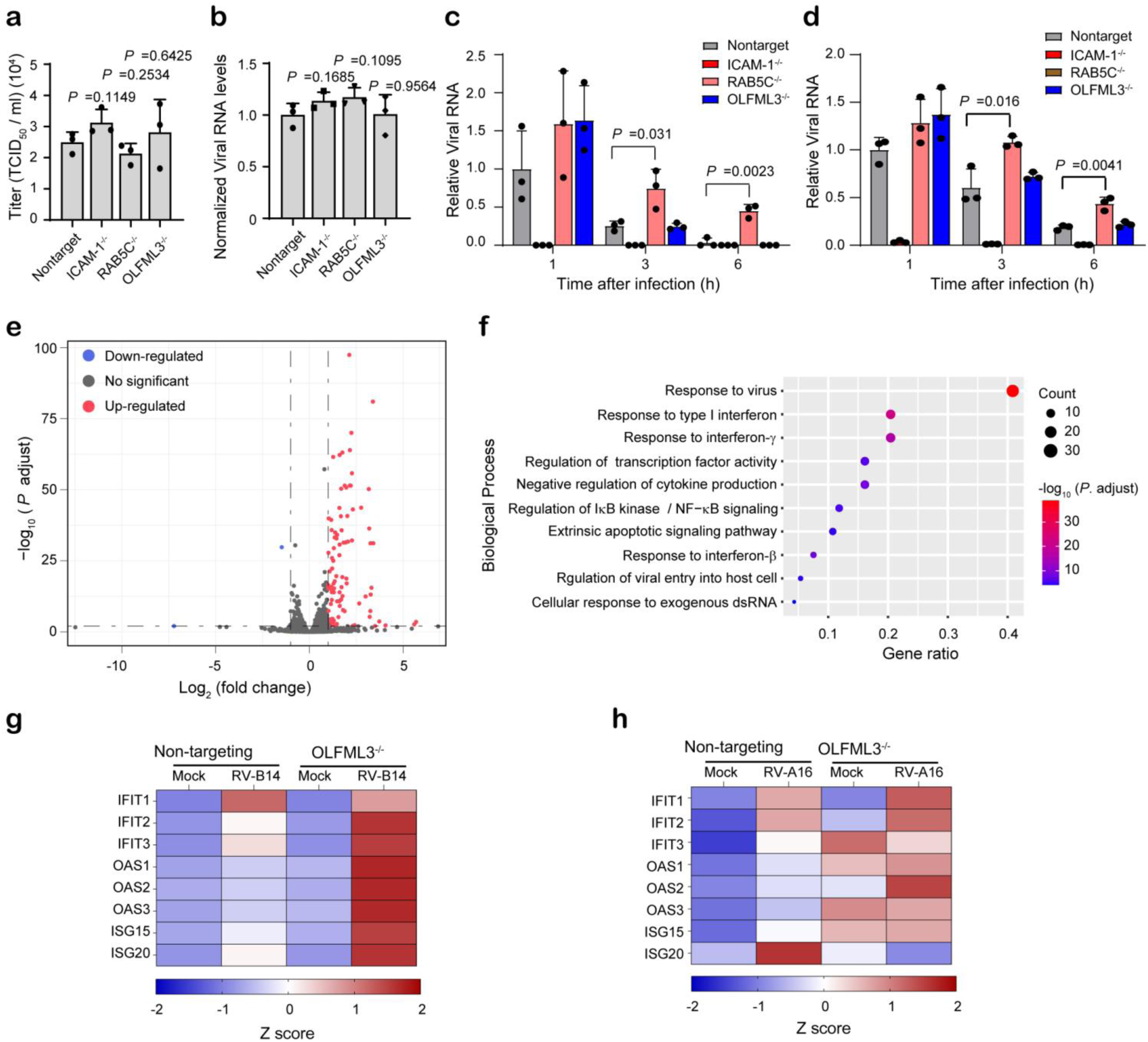
Dissection of the functions of RAB5C and OLFML3 in RV infection. **a-b** Viral loads in medium supernatant (**a**) and cell lysates (**b**) at 24 h after transfection of RV-A16 genome RNA. **c-d** Viral RNA in cell lysates of mock and knockout cells at 1, 3 and 6 h after infection with RV-B14 (**c**) and RV-A16 (**d**) at an MOI of 20 in the presence of 2 mM GuHCl. For **a-b**, significant difference between mock and test groups is determined using two-tailed Student’s *t* test. For **c-d**, significant difference between mock and RAB5C groups is determined using two-tailed Student’s *t* test. **e** Volcano plot showing differentially expressed genes (DEGs). RV infection-induced gene upregulation and downregulation are first calculated and the differentially upregulated or downregulated genes in mock and knockout cells are defined as DEGs. Cells are harvested and analysed at 24 h after infection of RV-B14 at an MOI of 2. **f** GO analyses of biological processes of DEGs identified in E. **g**-**h** Heat map showing RT-qPCR quantification of ISG expression in mock and OLFML3-/- cells at 24 h post infection of RV-B14 (**g**) and RV-A16 (**h**) at an MOI of 2. Gene expression is normalized to RPLP0.

### Identification of OLFML3 as a negative regulator of innate immune response

The above results showed that OLFML3 was a critical RV dependency factor but did not participate in RV attachment, entry or genome uncoating. This prompted us to elucidate the mechanism of action of OLFML3 in RV infection using transcriptome-wide RNA sequencing (RNA-Seq). RNA-Seq analyses were performed with wild-type and OLFML3^-/-^ cells before and after RV-B14 infection. The RNA-Seq data from three independent experimental replicates were collected and the replicates within each condition were found to have high degree of correlation (Additional file 1: Fig. S10a). Analyses of mapped read counts of RV-B14 genome revealed significant, transcriptome-wide inhibition of viral gene expression in OLFML3^-/-^ cells (Additional file 1: Fig. S10b), consistent with the above results of OLFML3 knockout-mediated inhibition of viral replication (Fig. 3). Analyses of differentially expressed genes (DEG) showed that removal of OLFML3 induced the upregulation of a broad range of genes upon RV-B14 challenge (Additional file 1: Fig. 4e and Fig. S10c). Gene ontology analyses of the biological processes and molecular function of DEGs revealed markedly changed innate immune response (Fig. 4f and fig. S10d). To validate the RNA-Seq results, we infected non-targeting sgRNA-transduced and OLFML3^-/-^ cells with RV-B14 and RV-A16 at an MOI of 2 for 24 h. RT-qPCR quantification uncovered a series of upregulated IFN-stimulating genes (ISGs) in OLFML3^-/-^ cells, but not in mock cells, upon RV infection (Fig. 4g-h). These results were consistent with the RNA-Seq analyses and strongly indicated that OLFML3 promoted RV infection in H1-Hela cells by antagonizing innate immune response.

### OLFML3 is a RV-inducible IFN suppressor

Although our results have suggested a possible role of OLFML3 in the innate immune response, the signaling pathway OLFML3 is involved in remains elusive. OLFML3 is secreted glycoprotein consisting of approximately 400 amino acids. OLFML3 belongs to the olfactomedin (OLF) superfamily and bears a C-terminal olfactomedin-like (OLFML) domain (Fig. 5a). In human, there are five OLFML members and the functions of OLFML1, OLFML2A, OLFML2B and OLFML4 have been illustrated [39]. However, the role of OLFML3 is poorly understood and there have been no studies reporting its functions with viral infection.

**Fig. 5.**
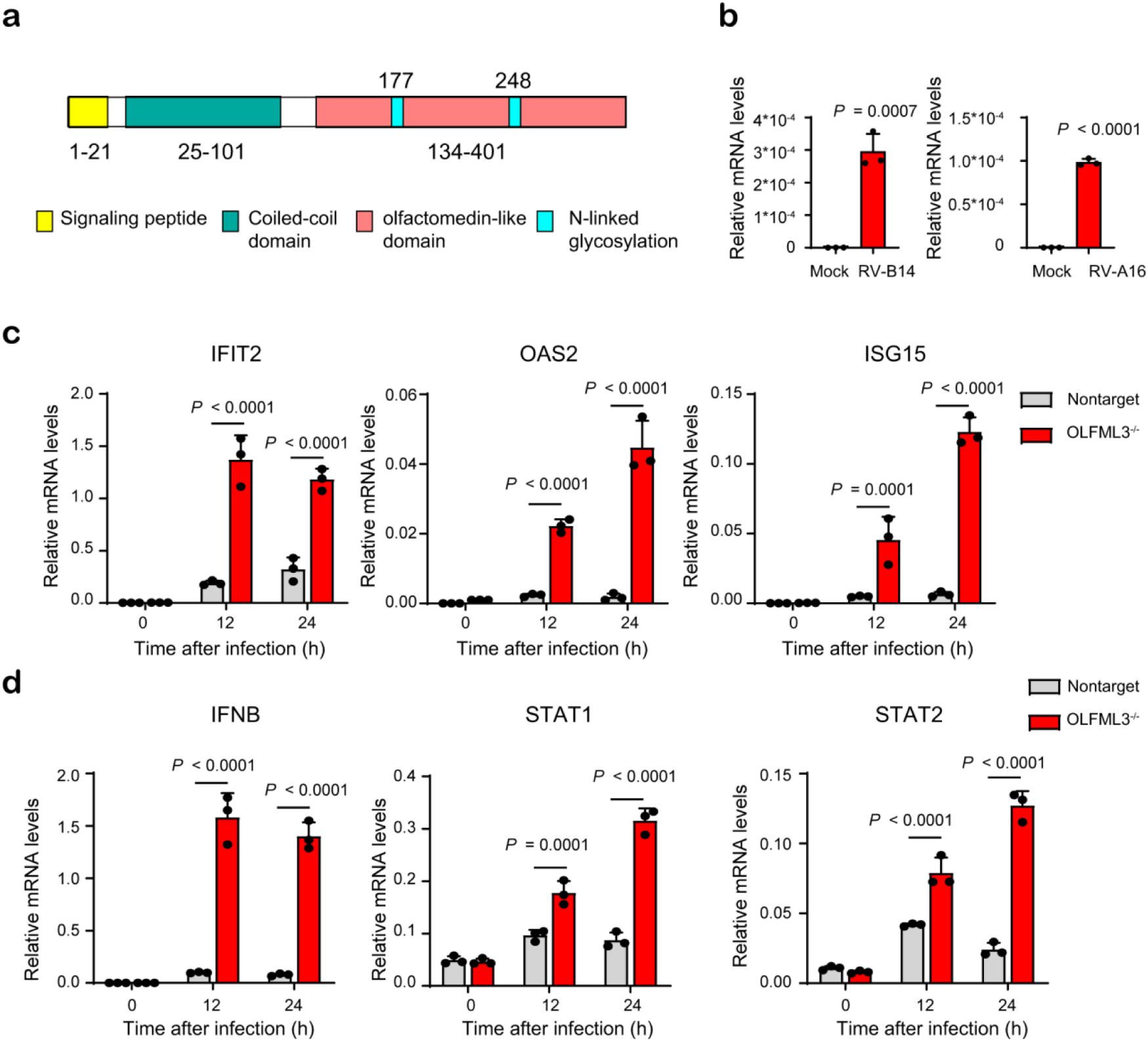
OLFML3 is a RV-inducible suppressor of type I IFN signaling during RV infection. **a** Structural organization of OLFML proteins. **b** RT-qPCR quantification of OLFML3 expression levels in H1-Hela cells in the absence or presence of RV-B14 or RV-A16. Samples are collected at 24 h post infection with a RV MOI of 2. **c** RT-qPCR quantification of IFIT2, OAS2 and ISG15 expression in WT and OLFML3^-/-^ cells at 0, 12 and 24 h after RV-B14 infection at an MOI of 2. **d** RT-qPCR quantification of IFNB, STAT1 and STAT2 expression in WT and OLFML3^-/-^ cells at 0, 12 and 24 h after RV-B14 infection at an MOI of 2. For **c** and **d,** gene expression is normalized to RPLP0 and significant difference was determined using two-way ANOVA with Sidak’s multiple comparisons test. Gene abbreviations are: IFNB, interferon β; ISG15, IFN-stimulating genes 15; IFIT2, interferon induced protein with tetratricopeptide repeats 2; OAS2, 2’-5’-oligoadenylate synthetase 1; SOCS3, suppressor of cytokine signaling 3.

To understand the function of OLFML3, we quantified its expression in H1-Hela. Interestingly, OLFML3 had very low mRNA expression under uninfected conditions, and RV-B14 and RV-A16 infection upregulated the expression of OLFML3 by more than 500 and 400 folds respectively (Fig. 5b). Importantly, in the presence of OLFML3, ISGs could not be efficiently activated by RV infection over a course of 12 h. By contrast, removal of OLFML3 allowed time-dependent upregulation of ISGs in response to RV infection (Fig. 5c). These observations are consistent with the above results (Fig. 4) and demonstrated that OLFML3 promoted RV infection (Fig. 3) by suppressing innate immune response. Moreover, IFN and STAT1/2 expression underwent minor changes in response to RV infection in the presence of OLFML3 while OLFML3 knockout sensitized IFN and STAT1/2 upregulation in response to RV infection (Fig. 5d), which was consistent with the RNA-Seq results (Fig. 4). Collectively, our results suggested that RV-induced OLFML3 activation and OLFML3-mediated inhibition of type I IFN signaling might be a critical mechanism for RV to escape the innate immune system.

### OLFML3 antagonizes type I IFN signaling in an SOCS3-dependent mechanism

Consistent with the RNA-Seq results (Fig. 4), OLFML3 knockout significantly reduced the expression of suppressor of cytokine signaling-3 (SOCS3) (Fig. 6a-b), a well characterized suppressor of IFN signaling [40], at 24 h post RV infection. SOCS3 knockdown using siRNA (Fig. 6b) prevented RV replication in H1-Hela cells (Fig. 6c). Interestingly, SOCS3 knockdown seemed more efficient than OLFML3 knockout in inhibiting viral replication (Fig. 6c). Meanwhile, SOCS3 knockdown and OLFML3 knockout did not have additive effects (Fig. 6c). These results suggested that OLFML3 and SOCS3 were in the same signaling pathway and that SOCS3 functioned as the predominating molecule or was downstream of OLFML3.

**Fig. 6.**
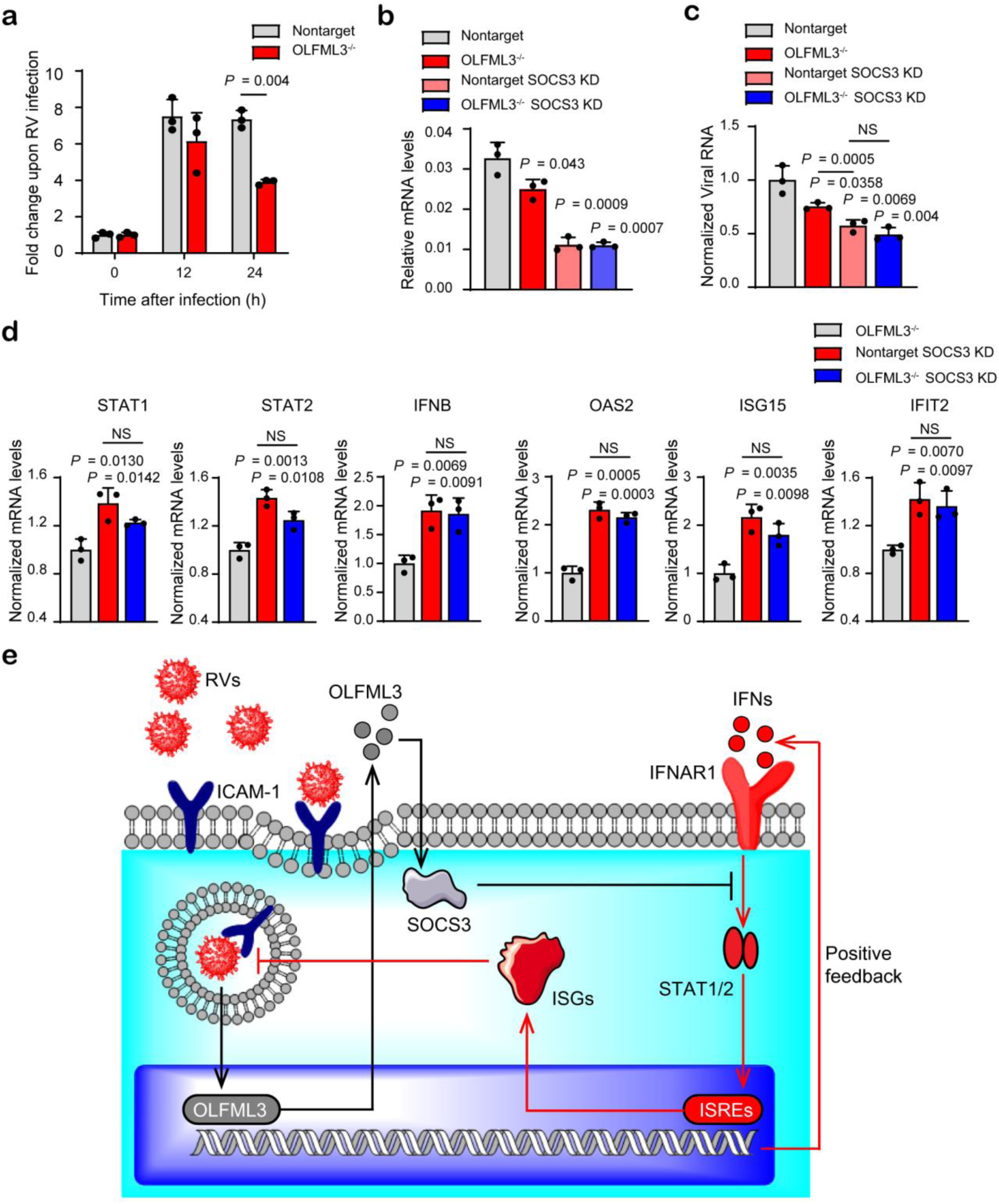
OLFML3 antagonizes type I IFN signaling in an SOCS3-dependent mechanism. **a** mRNA expression of SOCS3 in response to RV-B14 infection. Samples are collected at 0, 12 and 24 h after infection at an MOI of 2 and the fold change of SOCS3 expression before and after RV infection is shown. **b** RT-qPCR quantification of SOCS3 expression in mock or OLFML3^-/-^ cells in the absence and presence of SOCS3 siRNA. Samples are collected at 24 h post RV-B14 infection at an MOI of 2. **c** RT-qPCR quantification of viral RNA in cell lysate in mock or OLFML3^-/-^ cells in the absence and presence of SOCS3 siRNA. Samples are collected at 24 h post RV-B14 infection at an MOI of 2. Significant difference between non-target and each other group is determined unless indicated otherwise. **d** RT-qPCR quantification of STAT1, STAT2, IFNB, OAS2, ISG15 and IFIT2 expression in the absence and presence of SOCS3 siRNA. Samples are collected at 24 h post RV-B14 infection at an MOI of 2. Significant difference between OLFML^-/-^ and each other group is determined unless indicated otherwise. For **a**-**d,** gene expression is normalized to RPLP0 and significant difference is determined using Student’s *t* test. NS, no significance. **e** Schematic diagram illustrating OLFML3 and SOCS3-mediated inhibition of type I IFN signaling during RV infection. Gene abbreviations are: RVs, rhinoviruses; ICAM-1, intercellular adhesion molecule 1; OLFML3, olfactomedin like 3; SOCS3, suppressor of cytokine signaling 3; IFNs, interferons; IFNAR1, type I interferon receptor α chain; ISREs, interferon-stimulated response elements; ISGs, interferon-stimulating genes.

Next we analyzed the effects of SOCS3 knockdown on the expression of IFN signaling molecules. It was found that SOCS3 knockdown had more prominent impacts than OLFML3 knockout on activating IFN signaling molecules including STAT1/2, IFNB and ISGs. In consistency with the above viral infection, SOCS3 knockdown in OLFML3^-/-^ cells did not further enhance type I IFN signaling (Fig. 6d). These results consistently suggested that SOCS3 was a downstream molecule of OLFML3 that predominated the negative regulatory pathway of IFN signaling. Therefore, Our results collectively suggested that OLFML3 exerted its IFN-inhibiting activity through OLFML3-SOCS3-STAT1/2 axis (Fig. 6e).

## Discussion

The widespread RV infections and its remarkable phenotypic diversity have precluded the development of effective vaccines and antiviral therapeutics. This obstacle has rendered host-directed therapy (HDT) [41] an attractive option for treating RV infections, where host-virus interactions are interfered. HDT of RVs requires systemic dissection of cellular host factors supporting viral infection. Most importantly, as potential drug targets, the identified host factors should be readily accessible to antiviral therapeutics including macromolecular drugs. This prompted us to investigated surface protein-focused functional genomics approaches for uncovering druggable cell surface host factors of RVs. It is surprising to find that compared with genome-wide CRISPR screen, surfaceome screen is not only more efficient in identifying cell surface host factors but also exhibits an overall higher positive/success rate. Thus, our results have suggested that surfaceome CRISPR screen outperforms genome-wide screen in identifying viral host factors. We envision that this unique strategy could be rapidly adapted to other viruses to facilitate the identification of viral host factors.

The identified RV host factor RAB5C is a endosome-localized small GTPase and is deemed to involve in cellular trafficking [42]. It has been reported that RAB5C plays critical roles during the infection of flaviviruses including Zika virus and Dengue virus [35] as well as other viruses [43]. However, the roles of RAB5C during RV infection remains elusive. We found that RAB5C did not affect RV attachment or entry nor did it affect the replication of RV genome. Further analyses suggested that RAB5C might involve in the uncoating process of RV genome. Considering the reported cellular localization and function of RAB5C, our results indicate that RAB5C may be a key regulator for the endosomal release of RV genome.

Unlike RAB5C, OLFML3 does not participate in RV genome uncoating nor virus attachment or entry. OLFML3, also known as hOLF44, is one of the five OLFML proteins in human. OLFML1 is found to be associated with cell proliferation and autonomy in human cancer cells. OLFML2A and OLFML2B function as photomedin proteins. OLFML4 is known to have anti-inflammatory and anti-apoptotic activities. However, existing studies do not clearly define the biological functions of OLFML3. Some studies have suggested that OLFML3 may be involved in development and tumorigenesis [39] but none has reported its role in viral infection. In the present study, we found that OLFML3 promotes RV infection by acting as a type I IFN antagonist. The expression of OLFML3 is induced by RV infection, which may be an important mechanism for RV to escape from the innate immunity of host cells. We have shown that OLFML3-mediated inhibition of type I IFN signaling is dependent on SOCS3, a known suppressor of IFN signaling. Our study is the first to define OLFML3 as an IFN signaling inhibitor and to connect the functions of OLFML proteins to viral infections. Nevertheless, it is not completely unreasonable to expect the activities of OLFML3 in viral infections given the general roles of other OLFML proteins in inflammation and apoptosis and the signaling molecules these pathways share.

In future studies, it would be interesting to explore how OLFML3 initiates the negative regulatory pathway of IFN signaling. Particularly, it is important to investigate whether OLFML3 triggers the signaling through certain cellular receptors or whether OLFML3 directly interfere with the interactions between IFN and IFN receptors. In addition, it remains unclear how OLFML3 affects the expression of SOCS3. The fact the OLFML3 affected RV-activated SOCS3 expression at late time point (24 h after infection) suggested an indirect or delayed interactions between OLFML3 and SOCS3. Furthermore, it will be interesting to investigate whether OLFML3 functions as an antagonist of IFN signaling during the infection of other enteroviruses.

Finally, since OLFML3 is a secreted glycoprotein that can be readily accessed by drug molecules, it is worth developing small molecule or antibody therapeutics targeting OLFML3. On one hand, targeting OLFML3 with drug molecules can help elucidate its signaling pathway during RV infection. On the other hand, blocking OLFML3 with drug molecules can help assess the feasibility of OLFML3 as a drug target for treating RV infection. Importantly, because we have shown that OLFML3 does not function as attachment or entry receptor, simply blocking the solvent-exposed surface of OLFML3 may not be sufficient for therapeutics to have anti-RV effects. Thus, a screening platform may be needed to identify functional epitopes on OLFML3 for drug design and development.

## Conclusion

In the present study, we have shown that surfaceome CRISPR screen outperforms genome-wide screen in identifying RV host factors. Surfaceome screen has identified OLFML3 as a RV-inducible IFN suppressor that exerts the IFN-inhibiting activity through OLFML3-SOCS3-STAT1/2 axis. Our study has thus underscored surfaceome CRISPR screen for rapid dissection of host-pathogen interactions.

## Methods

### Design and construction of genome-wide and surfaceome CRISPR libraries

sgRNAs were designed to target to protein coding regions (NCBI CCDS data, released on 8-Sep-2016) [44] and optimized by two steps. First, off-target scores were calculated according to an established algorithm [30]. Second, on-target scores were calculated using Rule Set 2 [29]. sgRNAs were ranked by on-target scores and the top 12 sgRNAs with off-target scores of less than 20 were selected for each gene. If less than 12 sgRNAs were obtained, the cutoff of off-target scores was increased sequentially to 40, 60, 80 and 100 until 12 sgRNAs were obtained. The genome-wide CRISPR library was divided into three sub-libraries A, B and C according to sgRNA rank. Non-targeting sgRNAs were included in each library.

Pooled sgRNA oligonucleotides were synthesized as 76-mers by Custom Array (Bothell, WA, USA) and were amplified by PCR with NEBNext High Fidelity PCR Master Mix (New England BioLabs, NEB, Ipswich, MA, USA) using customized primers (Additional file 2: Table S1). The PCR products were purified using MinElute PCR purification kit (Qiagen, Dusseldorf, Germany). Lentiviral vector LentiCRISPR-v2 was digested with Esp3I (Thermo Fisher Scientific, Waltham, MA, USA) at 37 °C for 3 h and gel-purified using Gel Extraction kit (Omega, Norcross, GA, USA). Purified digestion products were ligated to Esp3I-treated LentiCRISPR-v2 using Gibson assembly kit (NEB) following manufacturer’s instructions. The ligation product was purified by isopropanol precipitation and then transformed into electrocompetent *Escherichia coli* (Lucigen, Middleton, WI, USA). Transformed cells were plated on to 15 cm Luria-Bertani (LB) agar plates supplemented with 50 μg/mL ampicillin (Sangon Biotech, Shanghai, China). Approximately 1-3 × 10^7^ colonies were collected for each library to ensure 500-fold coverage. Plasmid DNA was extracted as pooled libraries using NucleoBond Xtra Maxi EF kit (Macherey-Nagel, Duere, Germany) and stored at −20 ℃.

### Cell culture

H1-Hela cells were obtained from the American Type Culture Collection (ATCC). HEK293T cells were obtained from the Cell Bank of Shanghai Institutes for Biological Science (SIBS) and were validated by VivaCell Biosciences (Shanghai, China). H1-Hela and HEK293T were grown in Dulbecco’s modified Eagle’s medium (DMEM, Thermo) supplemented with 10% fetal bovine serum (FBS, Thermo) and 1% penicillin-streptomycin (Thermo) and maintained at 37 ℃ in a fully humidified incubator containing 5% CO_2._ All cells were confirmed by PCR to be free of mycoplasma contamination.

### LV production and transduction

To produce LVs with high titers, HEK293T cells were seeded on to 6-well plates with 2 × 10^6^ cells per well for single sgRNA, or 10 cm petri dishes with 10^7^ cells for library construction. At 24 h after seeding, HEK293T cells at a confluence of 70%-90% were transfected with LV packaging plasmid pMD2.G, envelope plasmid psPAX and transfer plasmid pLentiCRISPR-v2 that carries single or pooled sgRNAs using Lipofectamine 3000 (Thermo). At 6 h after transfection, the medium was replaced with fresh medium. The medium supernatant containing LVs was harvested at 48 h post transfection by centrifugation at 2,000 rpm for 10 min, filtrated through a 0.45-μm filter and stored at −80 ℃.

H1-Hela cells were transduced with LVs at an MOI of 0.3 using spinfection. Briefly, H1-Hela cells were washed with phosphate buffered saline (PBS) and incubated with LVs in serum-free DMEM under centrifugation at 2,000 rpm for 2 h. Upon completion of spinfection, LV-containing medium was removed and cells were incubated in DMEM supplemented with 10% FBS and 2 μg/mL puromycin (Thermo) for 2 to 3 days to purge empty cells containing no LVs or sgRNAs. For each library, cells of more than 500-fold coverage of the library size were collected.

### RV production and infection

The full-length cDNA clones of RV-A16 (pR16.11, Cat. No. VRMC-8) and RV-B14 (pWR3.26, Cat. No. VRMC-7) were obtained from ATCC. To produce infectious viral RNA, RV-A16 and RV-B14 plasmids were linearized by SacI (NEB) digestion and then *in vitro* transcribed using HiScribe T7 Transcription Kit (NEB). The RNA transcripts were extracted using Trizol (Thermo) and chloroform (Titan, Shanghai, China), followed by isopropanol precipitation. Viral RNA was transfected into H1-HeLa cells using Lipofectamine 3000 (Thermo) to generate infectious RV-A16 or RV-B14 particles. At 48 h post transfection, the supernatant containing RVs were collected for further infection on H1-Hela cells to produce RVs with higher titers. The aliquots of purified RVs were stored at −80 ℃. Virus titers were determined by the 50% tissue culture infectious dose (TCID_50_) assay.

For RV infection, H1-Hela cells were seeded on to 96- or 12-well plates with a density of 2×10^4^ or 1.5×10^4^ cells per well, respectively. Unless noted otherwise, at 24 h after seeding, cells were infected with RVs at an MOI of 2 for 1.5 h, washed with PBS for three times and then cultured in DMEM (Thermo) supplemented with 10% fetal bovine serum (FBS, Thermo) and 1% penicillin-streptomycin (Thermo) for 24 h.

### CRISPR screen using RV-B14

H1-Hela cells carrying genome-wide sub-library A or surfaceome library were seeded on to 15 cm petri dishes. Approximately 1.5 ×10^7^ cells were seeded to ensure more than 200-fold coverage of sgRNA. These cells were infected with RV-14 at an MOI of 1 for 48 h. At the end point of RV-14 challenge, the cells were washed with PBS and attached cells were collected from the plates. Two biological replicates were performed for mock and test groups. Genomic DNA of the harvested cells was extracted using phenol: chloroform: isoamyl alcohol (v/v/v, 25:24:1) and then purified using ethanol precipitation.

### Next-generation sequencing (NGS) analyses of sgRNA enrichment

Genome-integrated sgRNAs were amplified from extracted genomic DNA by PCR using the primers containing Illumina adaptor (Additional file 2: Table S1). The PCR product was gel-purified and then analyzed on Illumina HiSeq 3000 platform by Genewiz (Suzhou, Jiangsu, China). After removing the adaptors, the 20 bp sgRNA was mapped to the reference sgRNA libraries with one nucleotide mismatch allowed for each sgRNA. Gini index was calculated to analyze the distribution of sgRNAs. The raw read counts were subjected to MAGeCK analyses [45] to determine the enriched sgRNA and genes. Enrichment of sgRNAs and genes was analyzed using MAGeCK (v0.5.7) by comparing the read counts from the cells infected with RV-B14 with those from uninfected cells. A false discovery rate (FDR) of less than 0.01 was applied to identify significantly enriched sgRNAs or gene knockout.

### Generation of CRISPR-Cas9 knockout cells

The transfer plasmid pLentiCRISPR-v2 carrying single sgRNA for targeted gene knockout was constructed as described above. Briefly, forward and reverse oligonucleotides encoding the 20 bp sgRNA (Additional file 2: Table S2) were annealed to generate double-stranded DNA with overhang that matched the sticky ends of Esp3I (Thermo)-treated pLeniCRISPR-v2 vector. Annealed sgRNA sequence was ligated into digested pLeniCRISPR-v2 and then transformed into DH5α *E. coli* (Tsingke, Beijing, China).

The LVs carrying single sgRNA were packaged and transduced on to cells as described above. To evaluate the knockout efficiency, the genomic DNA of edited cells were extracted using Quick Extraction kit (Lucigen). Target sites carrying gene-edited sequences were PCR amplified using gene-specific primers (Additional file 2: Table S3). The knockout efficiency of each sgRNA was determined using T7E1 analysis. Single clones were obtained by limited dilution and genotyped by Sanger sequencing to determine the mutations at each allele. Knockout of target proteins was verified by western blotting (WB).

### Cell viability assay

H1-Hela cells were seeded into 96- or 24-well plates with a density of 5,000 or 50,000 cells per well respectively. At 24 h after seeding, cells were infected with RVs at an MOI of 1 for 1.5 h, washed with PBS for three times and cultured in fresh medium for 24 h unless noted otherwise. Cell counting Kit-8 (CCK-8, Dojindo, Kumamoto, Japan) was applied to determine cell viability according to manufacturer’s instructions. The absorbance at 450 nm was determined by Enspire multimode plate reader (PerkinElmer, Waltham, MA, USA).

### Immunofluorescence (IF) and fluorescence *in situ* hybridization (FISH)

IF staining of RV-B14 was performed using mouse anti-RV VP3 antibody (1:50, clone G47A, Thermo) and Alexa Fluor Plus 488 goat anti-mouse IgG (H+L) (1:1,000, A32723, Thermo). IF images were acquired and analyzed using Operatta high-content analysis system (PerkinElmer). At least 2,000 fluorescent cells were imaged and quantified for each replicate.

FISH was performed using an RNAscope Multiplex Fluorescent V2 Assay kit (Advanced Cell Diagnostics, Newark, USA) according to the manufacturer’s instructions. After fixation and pretreatment, RV-B14 RNA was detected using an RVB RNA probe (Advanced Cell Diagnostics, Cat. No. 447141) and TSA Plus Fluorescein (PerkinElmer, Cat. No. NEL741E001KT). FISH images were acquired and analyzed using a TissueFAXS 200 flow-type tissue quantitative analyser (TissueGnostics GmbH, Vienna, Austria). At least 5,000 cells in each replicate were included in analyses.

### Real-time quantitative PCR (RT-qPCR)

To determine viral loads, the medium supernatant or cell lysate containing RVs were harvested at 24 h after infection with RVs at an MOI of 2 unless noted otherwise. The total RNA from supernatant or cell lysate was purified using Trizol (Thermo) and chloroform (Titan), followed by purification using isopropanol precipitation. Purified viral RNA (vRNA) was reverse transcribed into cDNA using PrimeScript RT reagent Kit with gDNA Eraser (Takara Bio Inc., Shiga, Japan). The number of RV genome copy in the medium supernatant was determined using RT-qPCR with general or serotype-specific Taqman probe and primers (Additional file 2: Table S4) on Applied Biosystems Q6 Real-Time PCR cycler. The absolute viral titers were calculated based on a standard curve of RV genome with known TCID_50_ and the *R* square of curve-fitting was guaranteed to be more than 0.99. The mRNA levels of RVs and IFN-stimulating genes (ISGs) in cell lysate were determined using RT-qPCR with SYBR green dye (Thermo) and specific primers (Additional file 2: Table S5) on Applied Biosystems Q6 Real-Time PCR cycler. All SYBR Green primers were validated with dissociation curves. The expression of vRNA and host genes in cell lysate is normalized to ribosomal gene RPLP0 (36b4).

### Gene knockdown using siRNA

H1-Hela cells were seeded on to 6-well plates with a density of 5×10^5^ cells per well. At 24 h after seeding, cells were transfected with 100 pmol SOCS3 siRNA (Genepharma, Shanghai, China) (Additional file 2: Table S6) using 7.5 μL Lipofectamine 2000 (Thermo) for 6 h, washed with PBS and then cultured in fresh DMEM (Thermo) supplemented with 10% fetal bovine serum (FBS, Thermo). At 48 h post transfection, cells were infected with RV-B14 at an MOI of 2 for 1.5 h, washed with PBS for three times and then cultured in fresh medium for 24 h. The medium supernatant or cell lysate containing RVs were harvested and lysed for total RNA extraction, and then the mRNA levels of RVs, SOCS3, STAT1/2 and IFN-stimulating genes (ISGs) in cell lysate were determined using RT-qPCR as described above.

### Virus attachment and entry assays

H1-Hela cells were seeded in 12-well plate with a density of 200,000 cells per well and incubated overnight. For virus attachment assay, cells were incubated with RV-B14 or RV-A16 at an MOI of 20 in cold medium on ice for 60 min, washed by PBS for three times and then the total RNA was extracted using Trizol (Thermo). For virus entry assay, cells were incubated with RV-B14 or RV-A16 at an MOI of 20 in cold medium on ice for 60 min, washed by PBS for three times and then treated with pre-warmed medium for 40 min at 37 ℃. Then cells were washed with PBS for three times and treated with 0.25% trypsin for 2 min (Thermo, Cat. No. 25200072) to remove surface-bound viral particles. The internalized viral RNA was extracted using Trizol (Thermo) and viral loads were determined using RT-qPCR and FISH.

### Rescue experiments by overexpression

ICAM1, RAB5C and OLFML3 genes were codon-optimized for expression in human cells and synthesized by Genewiz. The 20 bp sgRNA-targeting sites and PAM sequences were mutated with silent mutations. Myc and FLAG tags were added to the C-terminus of these genes for WB detection. These genes were cloned into the EcoRI and XhoI sites of pCAGG plasmid that carries a separate mScarlet fluorescent protein as a transfection reporter.

### WB analysis

For WB analysis, cells were lysed with RIPA buffer (Beyotime Biotechnology, Beijing, China) on ice for 10 min. The total protein concentration in cell lysate was determined using the BCA Protein Assay Kit (Thermo). Cell lysate was mixed with SDS-PAGE loading buffer (Takara) containing 200 mM DTT, incubated at 95 °C for 10 min and resolved on NuPAGE 4-12% Bis-Tris gels (Thermo). Protein samples were transferred onto nitrocellulose membranes using an iBlot gel transfer system (Thermo). The following primary and secondary antibodies were used in WB including anti-ICAM1 rabbit antibody (Cell Signaling Technology, Cat. No. 4915S, Danvers, USA), anti-RAB5C rabbit antibody (Thermo, Cat. No. PA551932), anti-OLFML3 rabbit antibody (Thermo, Cat. No. PA531581), HRP-conjugated anti-rabbit IgG (CST, Cat. No. 7074S). Anti-β actin antibody conjugated with HRP (Abcam, Cat. No. ab49900, Cambridge, UK) was used as an internal control.

### RNA-Seq analysis of mock and OLFML3^-/-^ cells

Non-targeting sgRNA-treated mock cells and OLFML3^-/-^ cells were seeded on to 10 cm plates at a density of 1×10^6^ cells per plate. At 24 h after seeding, the cells were infected with RV-B14 at an MOI of 1 or with PBS as mock infection. After 24 h of RV challenge, the remaining cells were washed with PBS and collected by Trizol (Thermo) treatment. Three biological triplicates were prepared for each group. The whole-transcriptome RNA sequencing was performed by Genewiz.

RNA-Seq short reads were aligned to the human genome (GRCh37) using Hisat2 (v2.0.1) [46]. Gene expression was counted as the number of short reads fully or partially aligned to the annotated gene model using HTSeq (v0.6.1) [47]. Genotype (non-targeting sgRNA-transduced mock cells and OLFML3^-/-^ cells) and treatment conditions (mock and RV-B14 infection) were the factor variables in our RNA-Seq data. DESeq2 (v1.26.0) [48] was used to examine the difference between the response of mock and knockout cells to RV infection, which was captured by the interaction term. RV infection-induced gene upregulation and downregulation were calculated and the differentially upregulated or downregulated genes in mock and knockout cells were defined as DEGs. Significant DEGs were filtrated with an adjusted *P* value of less than 0.05 and a fold change value of more than 2.

GO enrichment analysis was performed using cluster Profiler (v3.14.3) by comparing DEGs to a list of all human genes [49]. Adjusted *P* value of less than 0.001 or 0.05 was set as the filter for biological process and molecular function terms respectively.

### Isolation of clinical RV strain

Nasopharyngeal swab samples were collected from the hospitalized patients bearing respiratory infection symptoms such as fever, cough, pharyngalgia and others. Nasopharyngeal swab samples were maintained in viral-transport medium and all samples were conserved on −80°C until analyses. The swab samples were diagnosed for respiratory viruses and the remaining samples were used for isolation of RVs. All procedures were complied with the Measures for the Ethical Review of Biomedical Research Involving Human Subjects issued by the National Health and Family Planning Commission of the People’s Republic of China. The Ruijin Hospital Ethics Committee, Shanghai Jiaotong University School of Medicine, approved the sample collection protocol with a permit number of Ruijin Hospital Ethics Committee 2018-48.

For isolation of RVs, the specimens were centrifuged at 2,000 rpm for 10 min and the supernatant was collected for RT-qPCR analysis using universal RV primers (Supplementary Table 4). The samples with high RV loads were selected for subsequent RV isolation in H1-Hela cells. H1-Hela cells were seeded on to 12-well plates at a cell density of 100,000 cells per well. Specimen supernatant (700 μL) was mixed with 300 μL of fresh medium and incubated with cells under centrifugation at 2,000 rpm for 2 h. Upon completion of spinfection, the supernatant was replaced with fresh medium and cells were incubated for another 2 days. Thereafter, medium supernatant and attached cells were collected and subjected to quick freeze-thaw cycles for three times to release viral particles. The medium supernatant or cell lysate from above was added to cells on 12-well plates. At 24 h after incubation, the supernatant was replaced with fresh medium and cells were incubated for 2 to 3 days before reaching 100% confluence. Isolation of RV strain was confirmed by CPEs and sequencing results.

### Statistical analyses

All data are the results from at least three biological replicates and are shown as mean ± SD unless noted otherwise. Statistical analyses and graphing were performed with GraphPad Prism 7.0. The *P* values were determined using two-tailed unpaired Student’s *t*-test unless otherwise noted.

## Supporting information

Additonal file 3

## Supplementary Information

**Additional file1: Fig S1-S10**

**Fig. S1.**
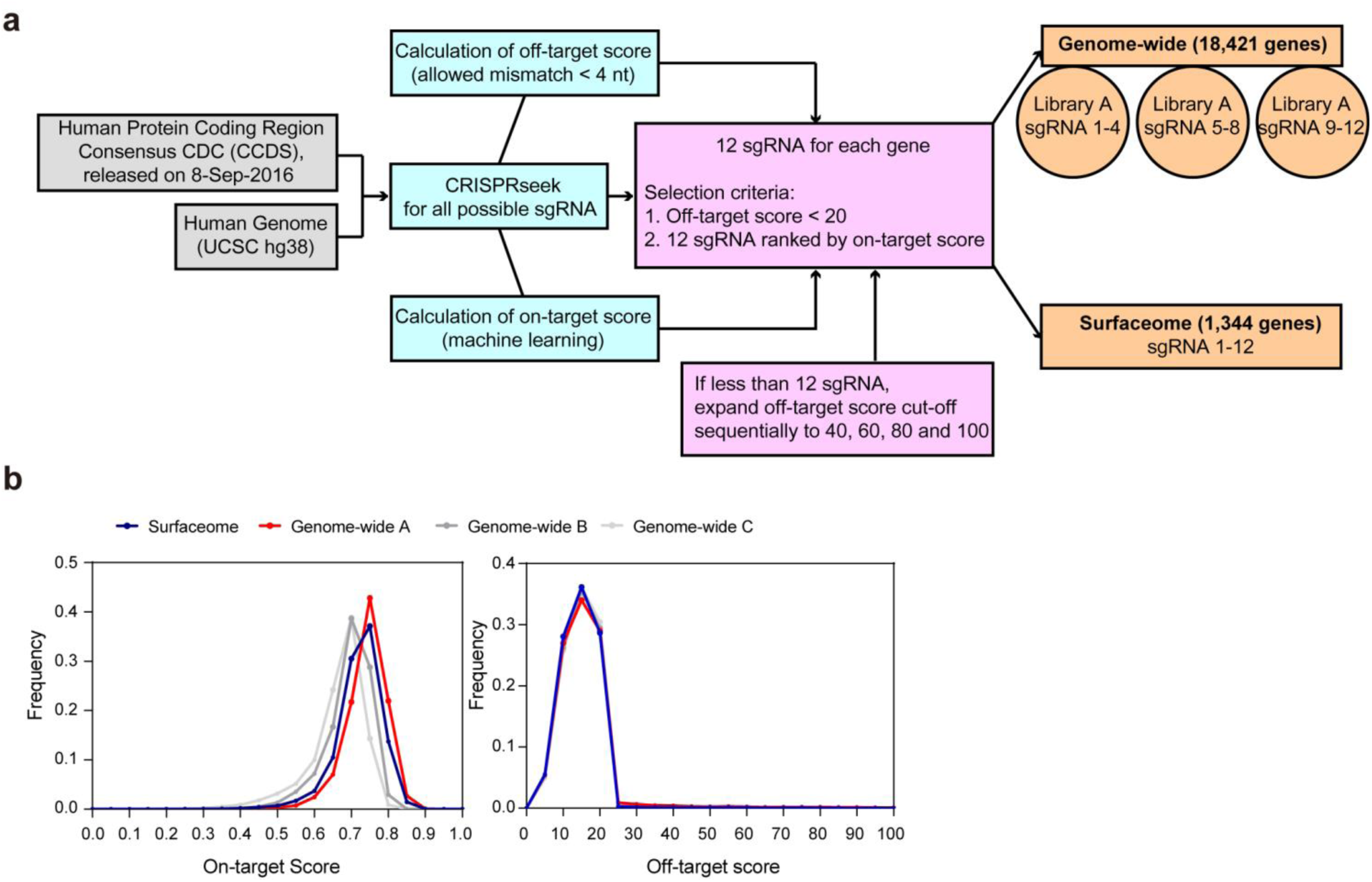
Construction of CRISPR genome-wide and surfaceome libraries. **a** Schematic illustration. **b** The distribution of sgRNA on-target and off-target scores in genome-wide sub-libraries A, B, C and surfaceome library.

**Fig. S2.**
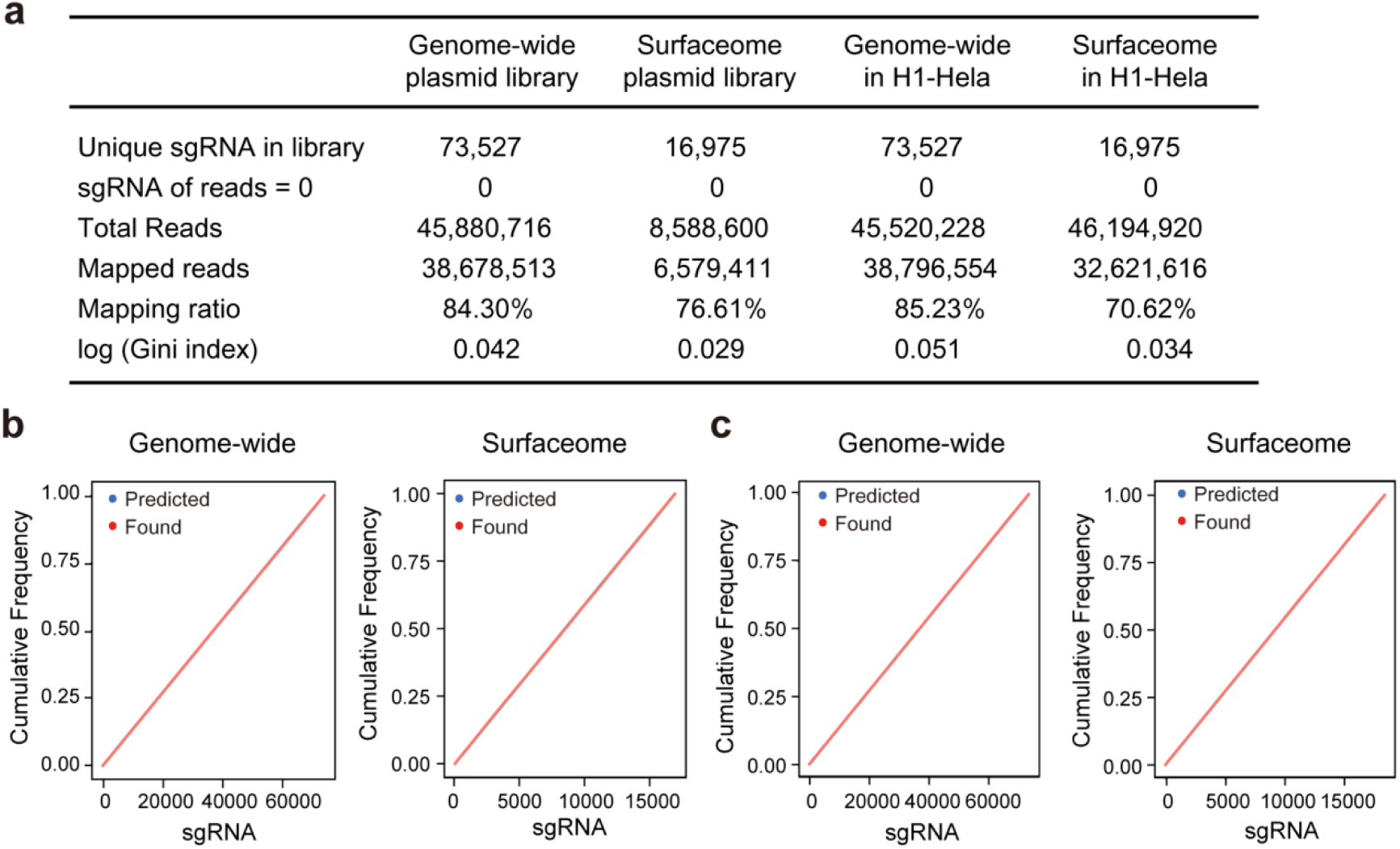
Quality analyses of constructed genome-wide and surfaceome CRISPR libraries. **a** Summary of next-generation sequencing results of sgRNA in the genome-wide and surfaceome plasmid and H1-Hela libraries. **b**-**c** Distribution of sgRNA in genome-wide and surfaceome libraries in pooled plasmids (**b**) and H1-Hela cells (**c**).

**Fig. S3.**
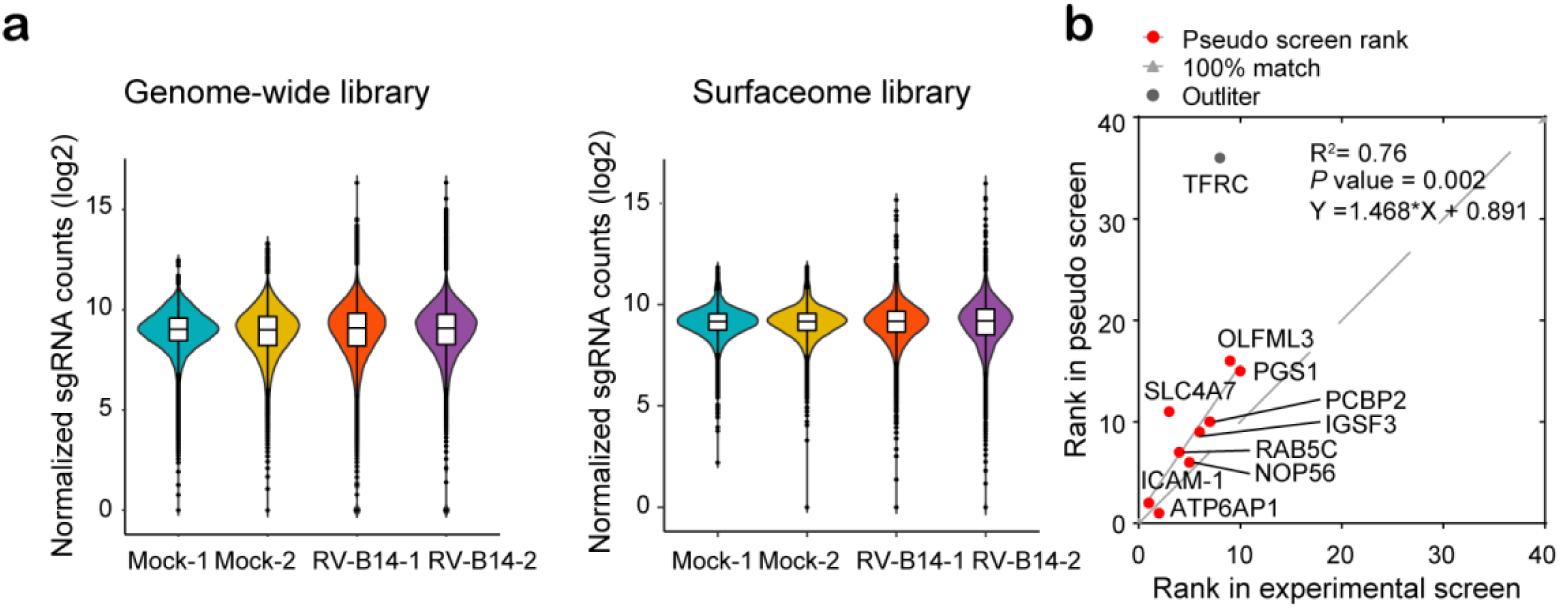
sgRNA distribution in surfaceome and genome-wide CRISPR libraries post RV-14 challenge. **a** Experimental results of sgRNA enrichment. **b** Correlation analyses of the top 10 hits from experimental surfaceome screen and their ranks in *in silico* (pseudo) surfaceome screen using sgRNA 1-4. The R square and *P* values of the correlation are calculated and shown.

**Fig. S4.**
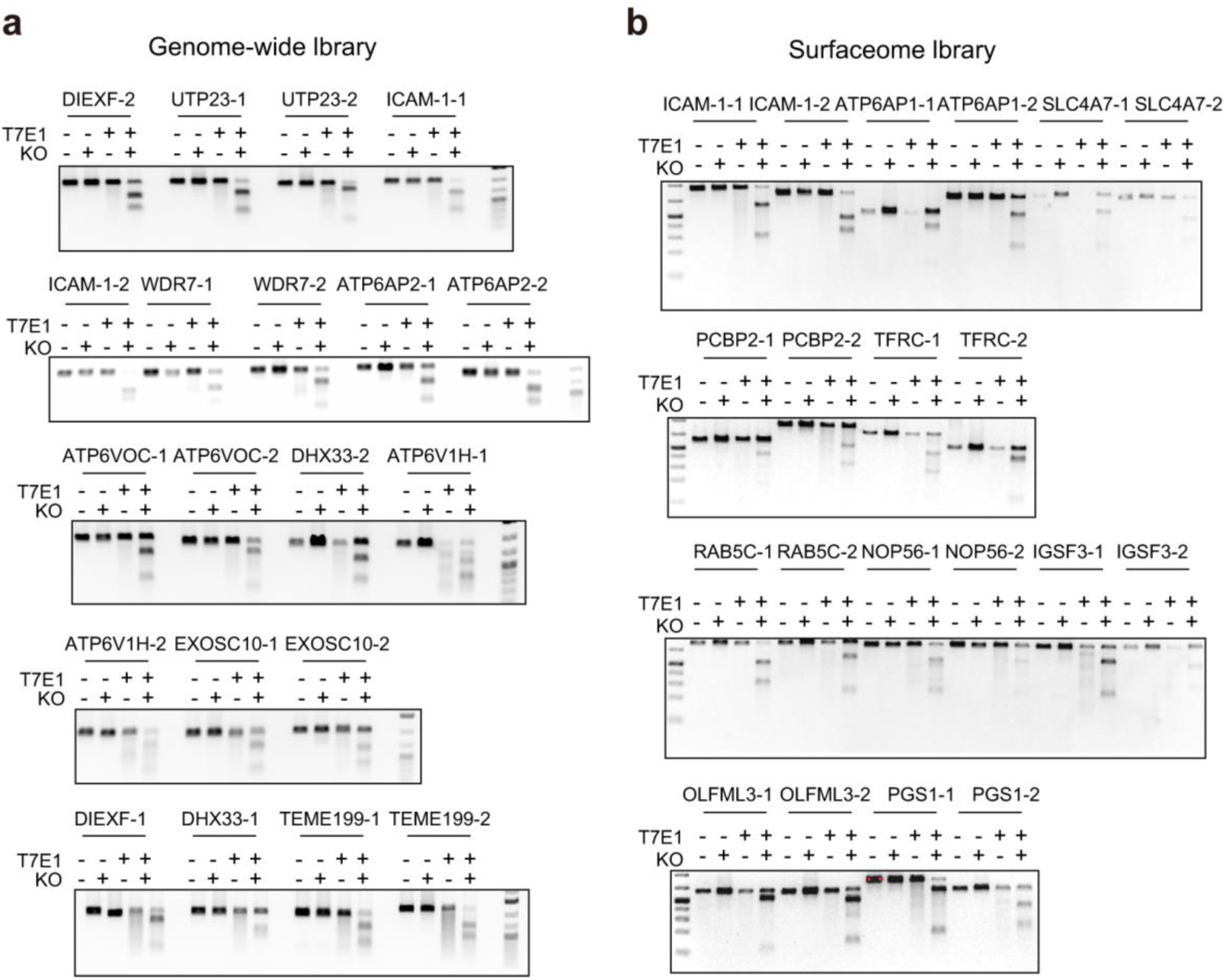
T7E1 analyses of the gene modification efficiency of sgRNA for candidate genes identified from surfaceome (a) and genome-wide screens (b).

**Fig. S5.**
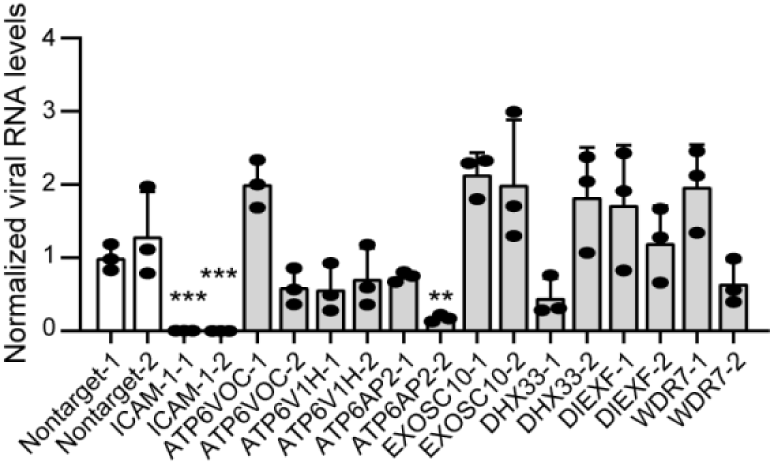
RT-qPCR quantification of viral loads in the lysate of knockout cells of the top 10 hits identified from genome-wide screen. Cell lysate is harvested at 24 h post RV-B14 infection at an MOI of 2. Viral RNA is normalized to RPLP0. The significant difference between mock and knockout cells is determined using two-tailed unpaired Student’s *t*-test, *P* = 0.0006 for ICAM-1-1 and ICAM-1-2, and *P* = 0.0015 for ATP6AP2-2.

**Fig. S6.**
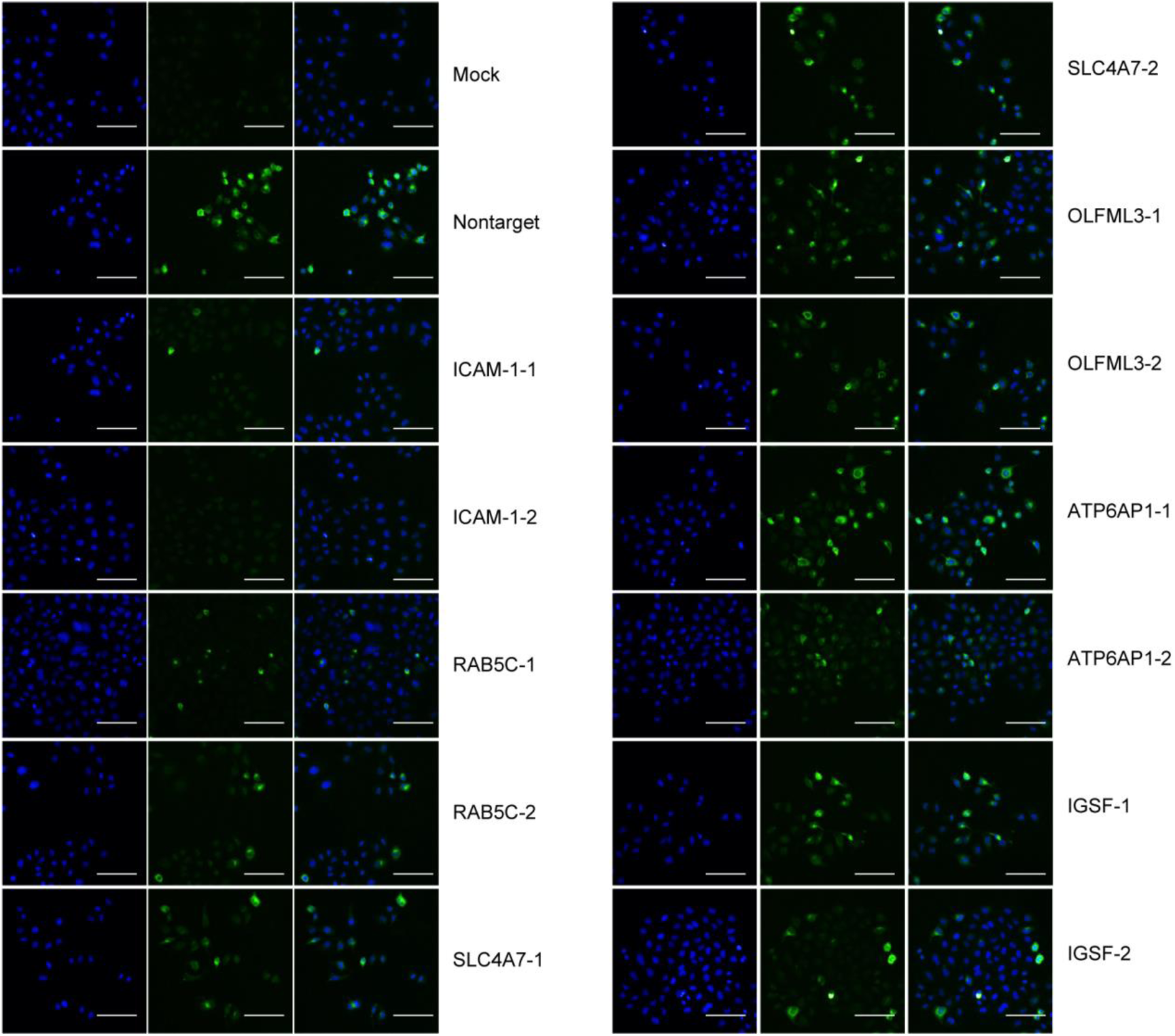
Representative IF images of knockout cells at 16 h post infection of RV-B14 at an MOI of 1. DAPI, blue; RV-B14 envelope protein, green. Scale bar, 100 μm.

**Fig. S7.**
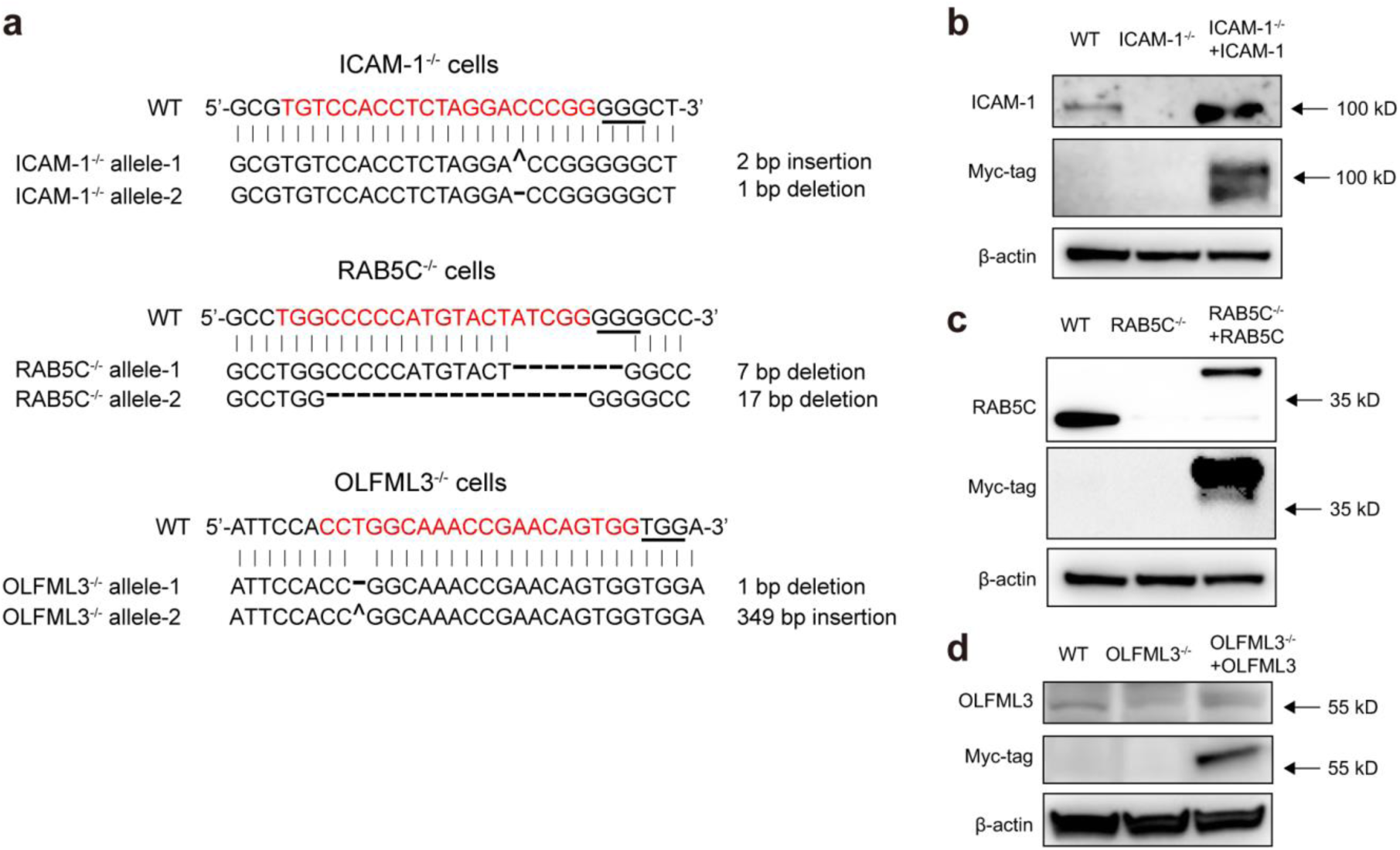
Validation of single clones of ICAM-1^-/-^, RAB5C^-/-^ and OLFML3^-/-^ H1-Hela cells. **a** Sanger sequencing analyses of mutated alleles. The 20-bp CRISPR-Cas9 target sequences are highlighted in red and protospacer adjacent motif (PAM) underlined. **b-d** Western blot analyses of ICAM-1 (**b**), RAB5C (**c**) and OLFML3 (**d**) expression in wide-type, knockout and rescued cells.

**Fig. S8.**
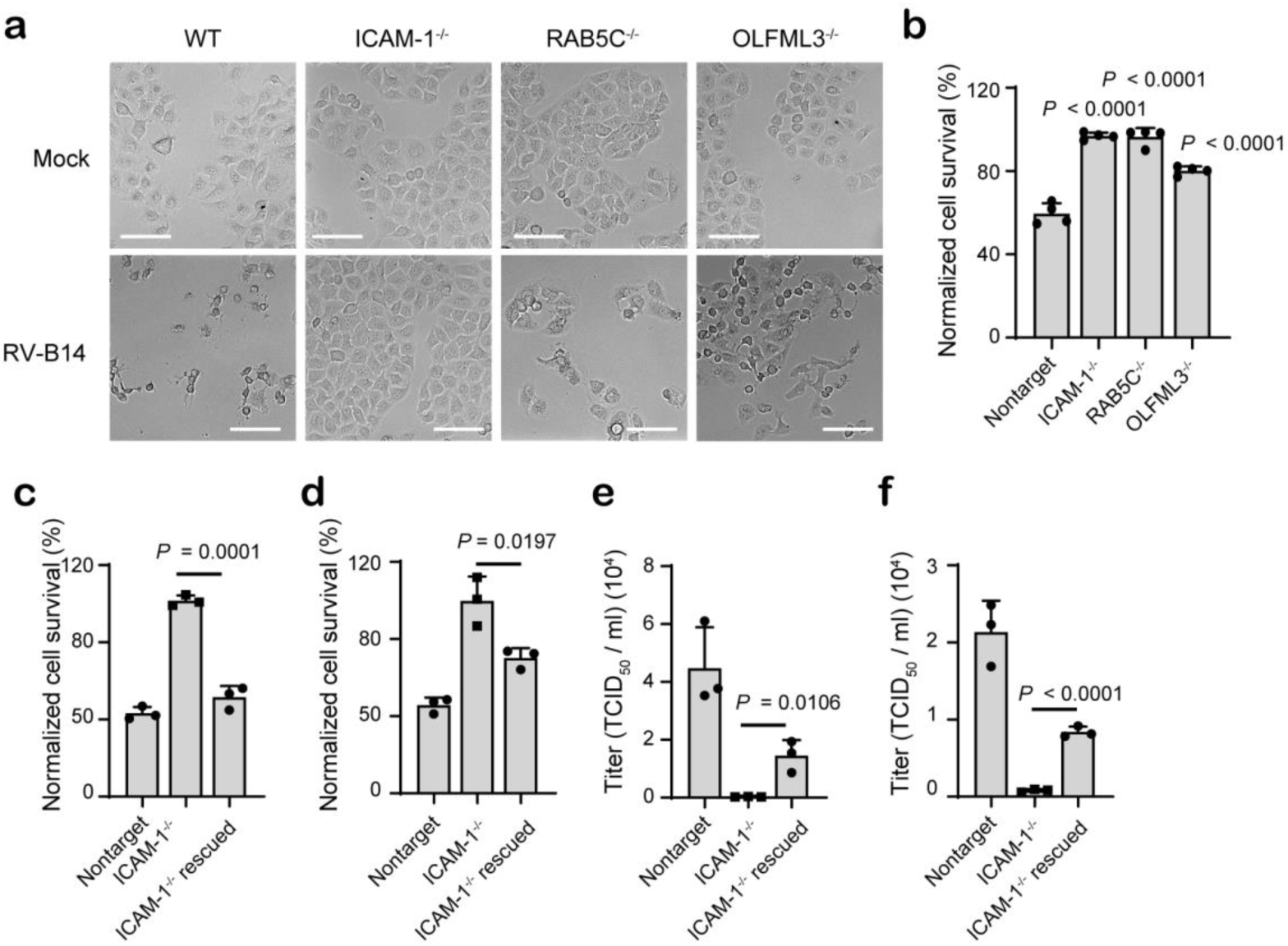
Validation of the effects of ICAM-1, RAB5C and OLFML3 on RV infection, related to Fig. 3. **a**-**b** Cell viability assay to determine the protective effects of ICAM-1, RAB5C and OLFML3 knockout against RV-B14 infection. Experiments are performed with an MOI of 2 and cell viability is determined at 24 h post infection. **a** Representative images. Scale bar, 100 μm. **b** Quantification of cell viability. Significant difference between test groups and non-targeting sgRNA group is determined using two-tailed Student’s *T* test and the *P* values are shown. **c**-**d** Cell viability assay to determine the effects of ICAM-1 overexpression on RV-B14 (**c**) or RV-A16 (**d**)-induced cell death in ICAM-1^-/-^ cells. **e-f** Rescued susceptibility of ICAM-1^-/-^ H1-Hela cells to RV-B14 (**e**) and RV-A16 (**f**) infection by ICAM-1 overexpression, as determined by viral loads in medium supernatant. Significant difference between knockout and rescued cells is determined using two-tailed unpaired Student’s *t* test.

**Fig. S9.**
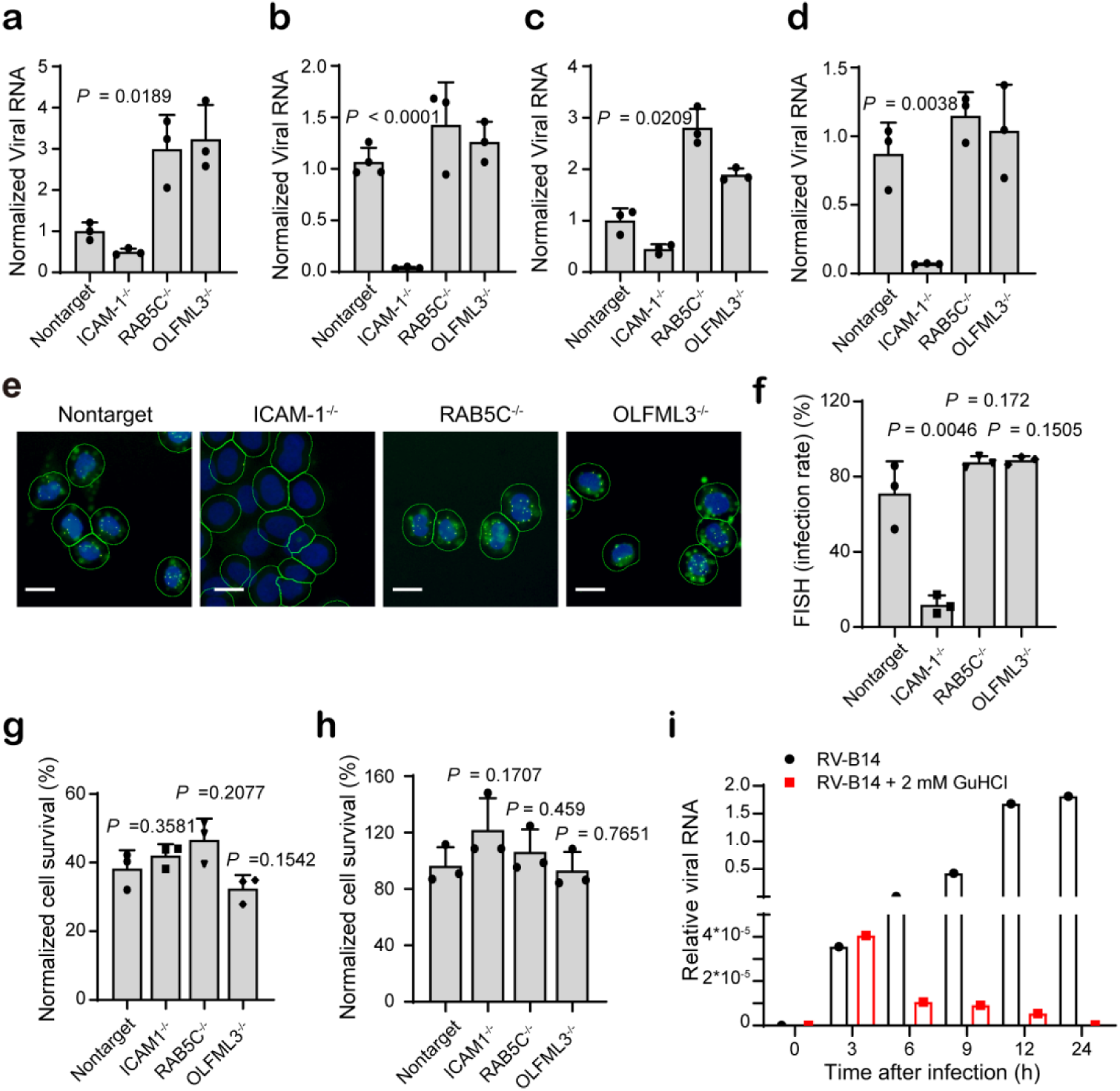
Dissection of the functions of RAB5C and OLFML3 in RV infection. **a-b** RV-B14 attachment (**a**) and entry (**b**) assays. **c**-**d** RV-A16 attachment (c) and entry (d) assays. For **a-d**, attached or internalized RV RNA is normalized to RPLP0. **e** Representative images. DAPI, blue; viral genome RNA, green dots; cell membrane, green lines. Scale bar, 20 μm. **f** Quantification of FISH experiments. The results are shown as mean ± SD (*n* = 3). In each replicate, 5,000 cells are analyzed. **g** Cell viability of mock and knockout cells at 24 h after transfection of RV-A16 genome RNA. **h** Cell viability at 6 h after treatment with 2 mM GuHCl. Significant difference between mock and knockout cells is determined using two-tailed unpaired Student’s *t* test. **i** Inhibition of the synthesis of RV-B14 viral RNA by treatment with 2 mM GuHCl. Cells are infected with RV-B14 at an MOI of 20 and viral RNA in cell lysates is determined and normalized to RPLP0.

**Fig. S10.**
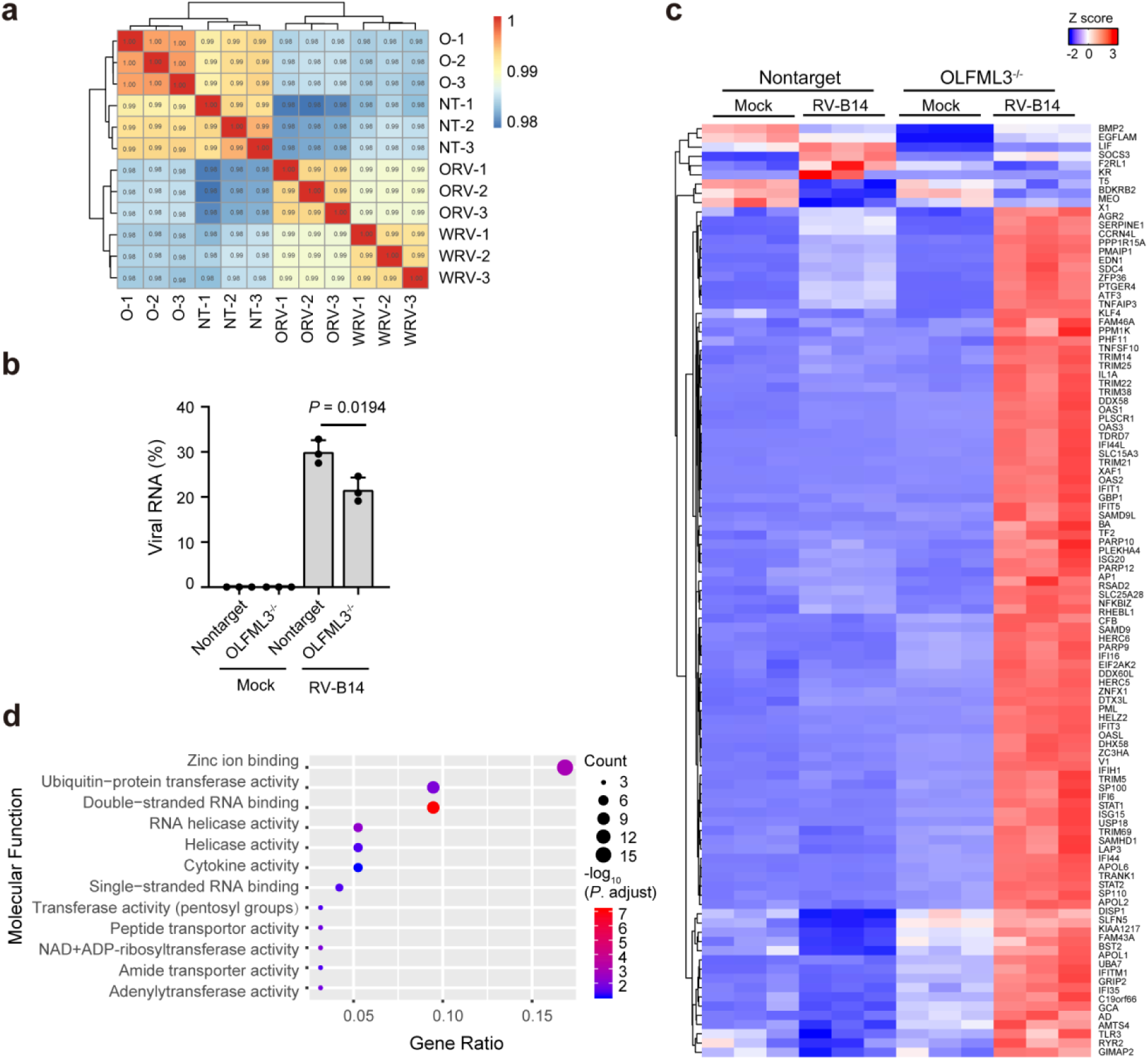
RNA-Seq analyses of the effects of OLFML3 on RV infection (related to Fig. 4). **a** Pearson correlation analyses of sequenced samples. The symbols are NT for non-targeting sgRNA-transduced cells, O for OLFML3^-/-^ cells and RV for RV-B14 infection. **b** Analyses of the effects of OLFML3 knockout on the transcriptomic expression of RV-B14. The significant difference of RV transcriptomic expression between mock and OLFML3^-/-^ cells is determined using two-tailed unpaired Student’s *t* test. **c** Heat map showing the differentially expressed genes with adjusted *P* values of less than 0.05and fold change of more than 2. Cells are collected for RNA-Seq analyses at 24 h after infection with RV-B14 at an MOI of 2. **d** GO analyses of the molecular functions of DEGs.

**Additional file 2: Table S1-S6**

**Table S1.**
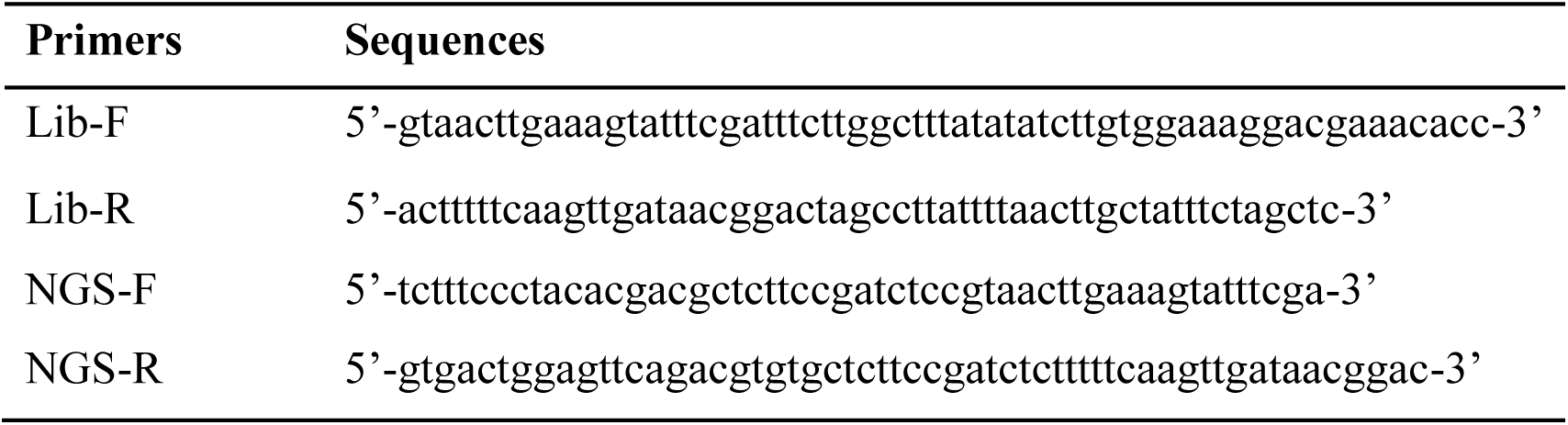
Primers for construction of CRISPR library and RNA-Seq analyses.

**Table S2.**
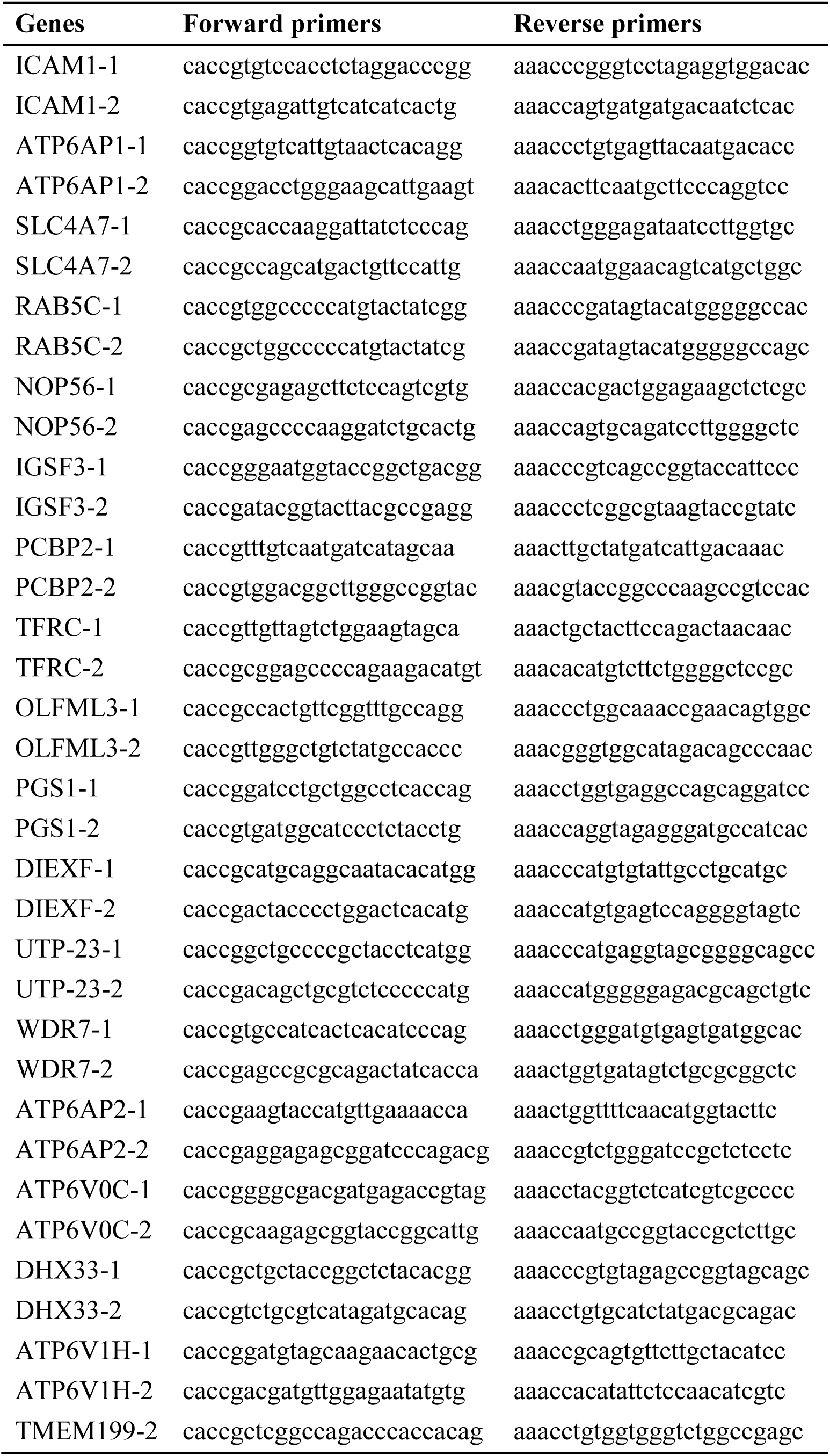
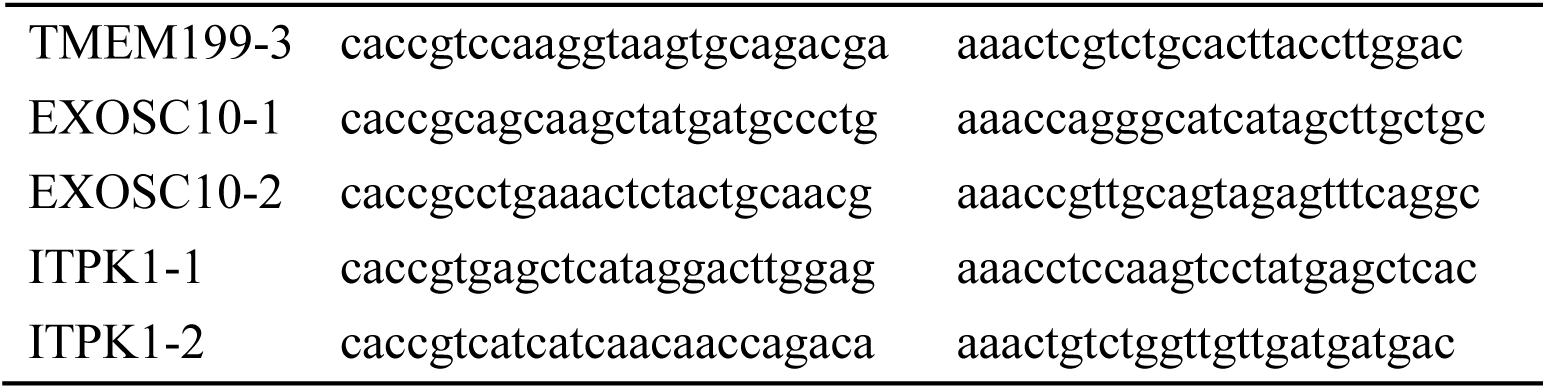
Primers for construction of sgRNA plasmid.

**Table S3.**
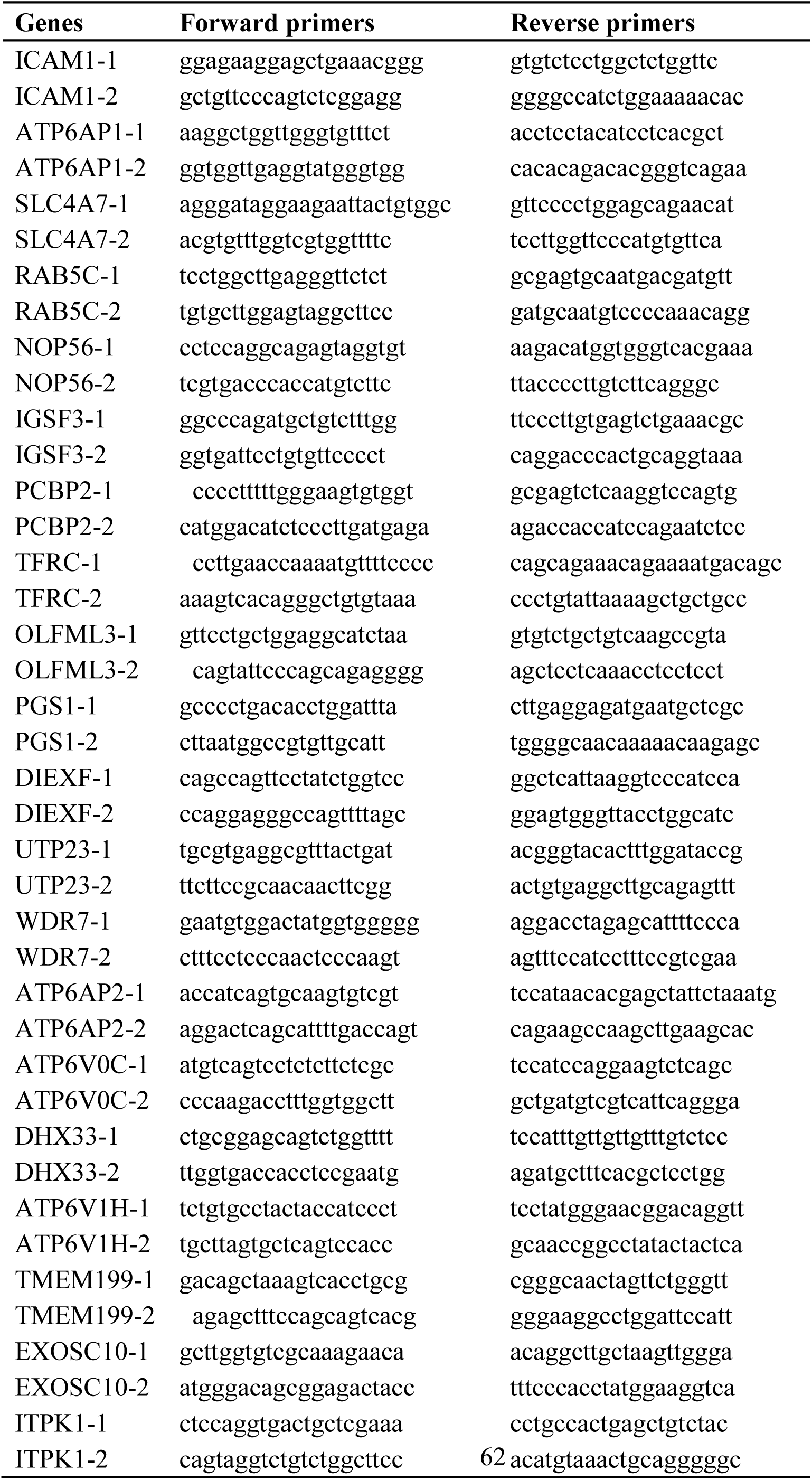
Primers for PCR amplification of sgRNA targeted sites for T7E1 analyses.

**Table S4.**
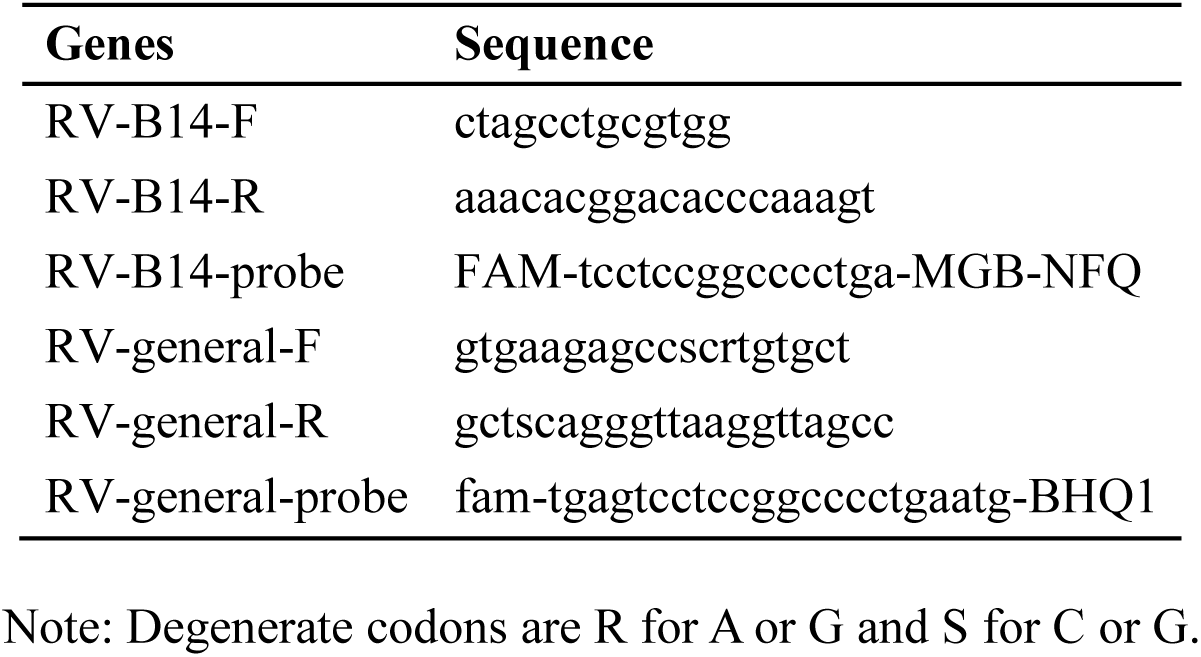
RV-B14-specific or RV universal primers for Taqman RT-qPCR.

**Table S5.**
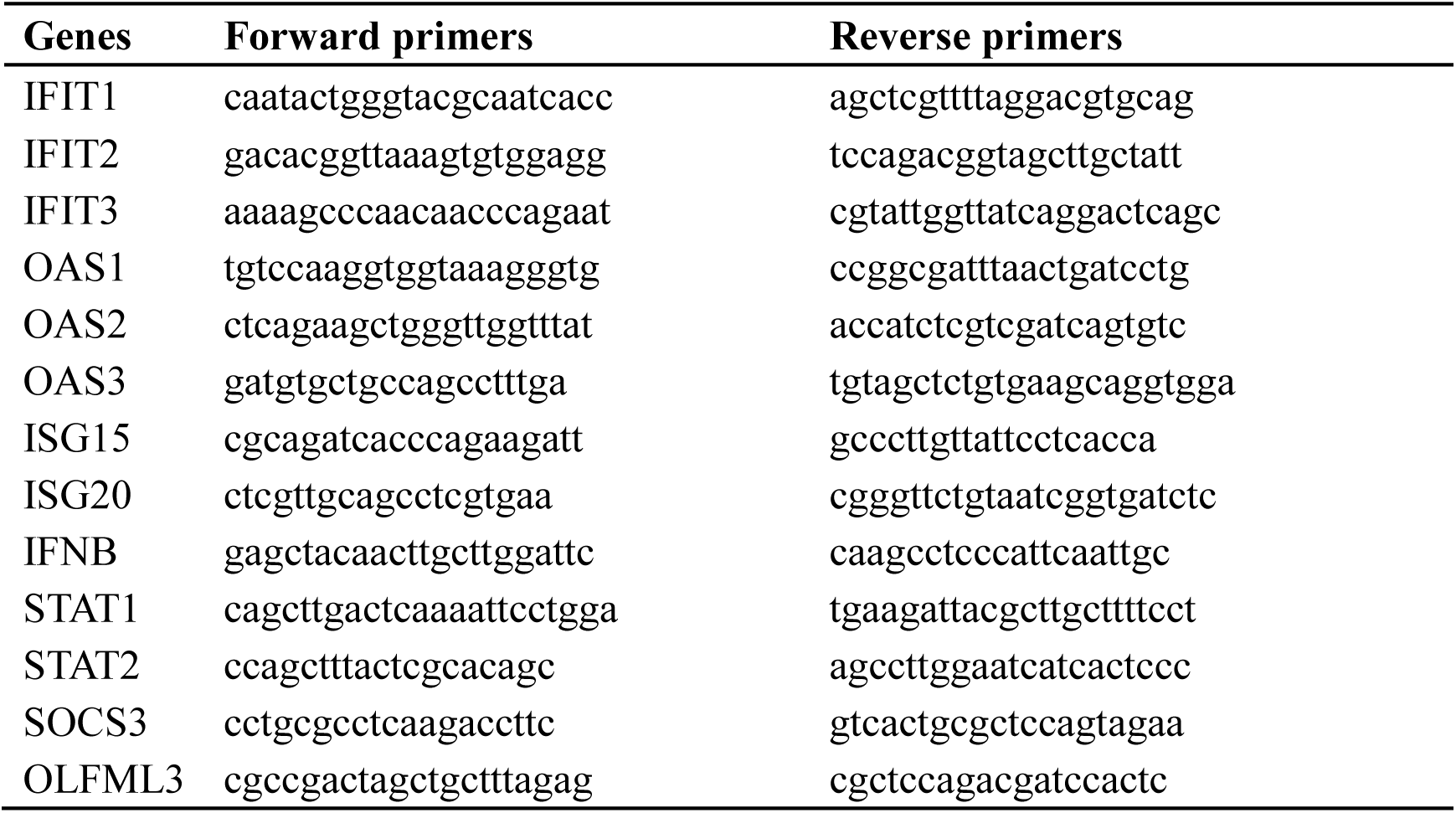
Primers for RT-qPCR.

**Table S6.**
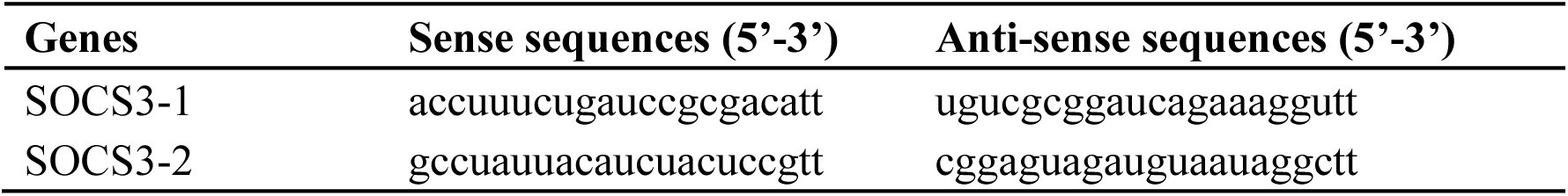
siRNA sequences.

**Additional file 3** Design of genome-wide CRISPR library A

**Additional file 4** Design of surfaceome CRISPR library

**Additional file 5** MAGeCK analysis of genome-wide screen

**Additional file 6** MAGeCK analysis of surfaceome screen

## Acknowledgements

We thank the High-throughput Screening (HTS) Platform at Shanghai Institute for Advanced Immunochemical Studies (SIAIS), ShanghaiTech University for the support of cell viability, IF and FISH experiments and Biomedical Big Data Platform for the design of sgRNA libraries, MAGeCK analyses and analyses of RNA-Seq data. We thank Professor Jincun Zhao (Guangzhou Medical University) for providing RV-B14 and RV-A16 genome plasmids.

## Author contributions

J.L. and J.Q. conceived this study and drafted the manuscript. J.L., J.Q., H.M. and Z.Z. designed the experiments. H.M., Z.Z., Y.X., W.L. and D.W. performed the experiments. W.W. and L.J. designed the sgRNA libraries, performed MAGeCK analyses and analyzed the RNA-Seq data. X.Z. and J.X. provided critical resources.

## Availability of data and materials

Additional information or materials supporting the findings in this study can be provided upon reasonable request to the corresponding author J. Liu.

## Ethics approval and consent to participate

Not applicable.

## Consent for publication

Not applicable.

## References

1. Doudna JA. The promise and challenge of therapeutic genome editing. Nature. 2020;578:229–236.

2. Doench JG. Am I ready for CRISPR? A user’s guide to genetic screens. Nat Rev Genet. 2018;19:67–80.

3. Puschnik AS, Majzoub K, Ooi YS, Carette JE. A CRISPR toolbox to study virus-host interactions. Nat Rev Microbiol. 2017;15:351–364.

4. Orchard RC, Wilen CB, Doench JG, Baldridge MT, McCune BT, Lee YC, et al. Discovery of a proteinaceous cellular receptor for a norovirus. Science. 2016;353:933–936.

5. Park RJ, Wang T, Koundakjian D, Hultquist JF, Lamothe-Molina P, Monel B, et al. A genome-wide CRISPR screen identifies a restricted set of HIV host dependency factors. Nat Genet. 2017;49:193–203.

6. Zhang R, Miner JJ, Gorman MJ, Rausch K, Ramage H, White JP, et al. A CRISPR screen defines a signal peptide processing pathway required by flaviviruses. Nature. 2016;535:164–168.

7. Marceau CD, Puschnik AS, Majzoub K, Ooi YS, Brewer SM, Fuchs G, et al. Genetic dissection of Flaviviridae host factors through genome-scale CRISPR screens. Nature. 2016;535:159–163.

8. Li B, Clohisey SM, Chia BS, Wang B, Cui A, Eisenhaure T, et al. Genome-wide CRISPR screen identifies host dependency factors for influenza A virus infection. Nat Commun. 2020;11:164.

9. Han J, Perez JT, Chen C, Li Y, Benitez A, Kandasamy M, et al. Genome-wide CRISPR/Cas9 screen identifies host factors essential for influenza virus replication. Cell Rep. 2018;23:596–607.

10. Staring J, von Castelmur E, Blomen VA, van den Hengel LG, Brockmann M, Baggen J, et al. PLA2G16 represents a switch between entry and clearance of Picornaviridae. Nature. 2017;541:412–416.

11. Zhao X, Zhang G, Liu S, Chen X, Peng R, Dai L, et al. Human neonatal Fc receptor is the cellular uncoating receptor for enterovirus B. Cell. 2019;177:1553–1565 e1516.

12. Diep J, Ooi YS, Wilkinson AW, Peters CE, Foy E, Johnson JR, et al. Enterovirus pathogenesis requires the host methyltransferase SETD3. Nat Microbiol. 2019;4:2523–2537.

13. Zhang R, Kim AS, Fox JM, Nair S, Basore K, Klimstra WB, et al. Mxra8 is a receptor for multiple arthritogenic alphaviruses. Nature. 2018;557:570–574.

14. Meertens L, Hafirassou ML, Couderc T, Bonnet-Madin L, Kril V, Kummerer BM, et al. FHL1 is a major host factor for chikungunya virus infection. Nature. 2019;574:259–263.

15. Jacobs SE, Lamson DM, St George K, Walsh TJ. Human rhinoviruses. Clin Microbiol Rev. 2013;26:135–162.

16. Castillo JR, Peters SP, Busse WW. Asthma Exacerbations: Pathogenesis, Prevention, and Treatment. J Allergy Clin Immunol Pract. 2017;5:918–927.

17. Seemungal T, Harper-Owen R, Bhowmik A, Moric I, Sanderson G, Message S, et al. Respiratory viruses, symptoms, and inflammatory markers in acute exacerbations and stable chronic obstructive pulmonary disease. American Journal of Respiratory and Critical Care Medicine. 2001;164:1618–1623.

18. Perreira JM, Meraner P, Brass AL. Functional genomic strategies for elucidating human-virus interactions: will CRISPR knockout RNAi and haploid cells? Adv Virus Res. 2016;94:1–51.

19. Perreira JM, Aker AM, Savidis G, Chin CR, McDougall WM, Portmann JM, et al. RNASEK is a V-ATPase-associated factor required for endocytosis and the replication of rhinovirus, influenza A virus, and Dengue virus. Cell Rep. 2015;12:850–863.

20. Haapaniemi E, Botla S, Persson J, Schmierer B, Taipale J. CRISPR-Cas9 genome editing induces a p53-mediated DNA damage response. Nat Med. 2018;24:927–930.

21. Wang C, Wang H, Lieftink C, du Chatinier A, Gao D, Jin G, et al. CDK12 inhibition mediates DNA damage and is synergistic with sorafenib treatment in hepatocellular carcinoma. Gut. 2020;69:727–736.

22. Tian R, Gachechiladze MA, Ludwig CH, Laurie MT, Hong JY, Nathaniel D, et al. CRISPR interference-based platform for multimodal genetic screens in human iPSC-derived neurons. Neuron. 2019;104:239–255 e212.

23. Li F, Huang Q, Luster TA, Hu H, Zhang H, Ng WL, et al. *In vivo* epigenetic CRISPR screen identifies Asf1a as an immunotherapeutic target in Kras-mutant lung adenocarcinoma. Cancer Discov. 2020;10:270–287.

24. Xu G, Chhangawala S, Cocco E, Razavi P, Cai Y, Otto JE, et al. ARID1A determines luminal identity and therapeutic response in estrogen-receptor-positive breast cancer. Nat Genet. 2020;52:198–207.

25. Inoue D, Chew GL, Liu B, Michel BC, Pangallo J, D’Avino AR, et al. Spliceosomal disruption of the non-canonical BAF complex in cancer. Nature. 2019;574:432–436.

26. Bersuker K, Hendricks JM, Li Z, Magtanong L, Ford B, Tang PH, et al. The CoQ oxidoreductase FSP1 acts parallel to GPX4 to inhibit ferroptosis. Nature. 2019;575:688–692.

27. Chong ZS, Ohnishi S, Yusa K, Wright GJ. Pooled extracellular receptor-ligand interaction screening using CRISPR activation. Genome Biol. 2018;19:205.

28. Ye L, Park JJ, Dong MB, Yang Q, Chow RD, Peng L, et al. *In vivo* CRISPR screening in CD8 T cells with AAV-Sleeping Beauty hybrid vectors identifies membrane targets for improving immunotherapy for glioblastoma. Nat Biotechnol. 2019;37:1302–1313.

29. Doench JG, Fusi N, Sullender M, Hegde M, Vaimberg EW, Donovan KF, et al. Optimized sgRNA design to maximize activity and minimize off-target effects of CRISPR-Cas9. Nat Biotechnol. 2016;34:184–191.

30. Hsu PD, Scott DA, Weinstein JA, Ran FA, Konermann S, Agarwala V, et al. DNA targeting specificity of RNA-guided Cas9 nucleases. Nat Biotechnol. 2013;31:827–832.

31. Bausch-Fluck D, Hofmann A, Bock T, Frei AP, Cerciello F, Jacobs A, et al. A mass spectrometric-derived cell surface protein atlas. PLoS One. 2015;10:e0121314.

32. Xie J, Yea K, Zhang H, Moldt B, He L, Zhu J, et al. Prevention of cell death by antibodies selected from intracellular combinatorial libraries. Chem Biol. 2014;21:274–283.

33. Li W, Koster J, Xu H, Chen CH, Xiao T, Liu JS, et al. Quality control, modeling, and visualization of CRISPR screens with MAGeCK-VISPR. Genome Biol. 2015;16:281.

34. Bella J, Kolatkar PR, Marlor CW, Greve JM, Rossmann MG. The structure of the two amino-terminal domains of human ICAM-1 suggests how it functions as a rhinovirus receptor and as an LFA-1 integrin ligand. Proc Natl Acad Sci U S A. 1998;95:4140–4145.

35. Savidis G, McDougall WM, Meraner P, Perreira JM, Portmann JM, Trincucci G, et al. Identification of Zika virus and Dengue virus dependency factors using functional genomics. Cell Rep. 2016;16:232–246.

36. Chen P, Hsu WH, Chang A, Tan Z, Lan Z, Zhou A, et al. Circadian regulator CLOCK recruits immune-suppressive microglia into the GBM tumor microenvironment. Cancer Discov. 2020;10:371–381.

37. Staunton DE, Merluzzi VJ, Rothlein R, Barton R, Marlin SD, Springer TA. A cell adhesion molecule, ICAM-1, is the major surface receptor for rhinoviruses. Cell. 1989;56:849–853.

38. Greve JM, Davis G, Meyer AM, Forte CP, Yost SC, Marlor CW, et al. The major human rhinovirus receptor is ICAM-1. Cell. 1989;56:839–847.

39. Jin Y, Li JL. Olfactomedin-like 3: possible functions in embryonic development and tumorigenesis. Chin Med J. 2019;132:1733–1738.

40. Rothlin CV, Ghosh S, Zuniga EI, Oldstone MB, Lemke G. TAM receptors are pleiotropic inhibitors of the innate immune response. Cell. 2007;131:1124–1136.

41. Schwegmann A, Brombacher F. Host-directed drug targeting of factors hijacked by pathogens. Sci Signal. 2008;1:re8.

42. Barbera S, Nardi F, Elia I, Realini G, Lugano R, Santucci A, et al. The small GTPase Rab5c is a key regulator of trafficking of the CD93/Multimerin-2/beta1 integrin complex in endothelial cell adhesion and migration. Cell Commun Signal. 2019;17:55.

43. Ziegler CM, Bruce EA, Kelly JA, King BR, Botten JW. The use of novel epitope-tagged arenaviruses reveals that Rab5c-positive endosomal membranes are targeted by the LCMV matrix protein. J Gen Virol. 2018;99:187–193.

44. Pruitt KD, Harrow J, Harte RA, Wallin C, Diekhans M, Maglott DR, et al. The consensus coding sequence (CCDS) project: Identifying a common protein-coding gene set for the human and mouse genomes. Genome Res. 2009;19:1316–1323.

45. Li W, Xu H, Xiao T, Cong L, Love MI, Zhang F, et al. MAGeCK enables robust identification of essential genes from genome-scale CRISPR/Cas9 knockout screens. Genome Biol. 2014;15:554.

46. Kim D, Langmead B, Salzberg SL. HISAT: a fast spliced aligner with low memory requirements. Nat Methods. 2015;12:357–360.

47. Anders S, Pyl PT, Huber W. HTSeq--a Python framework to work with high-throughput sequencing data. Bioinformatics. 2015;31:166–169.

48. Love MI, Huber W, Anders S. Moderated estimation of fold change and dispersion for RNA-seq data with DESeq2. Genome Biol. 2014;15:550.

49. Yu G, Wang LG, Han Y, He QY. clusterProfiler: an R package for comparing biological themes among gene clusters. OMICS. 2012;16:284–287.

50. Kumar S, Stecher G, Li M, Knyaz C, Tamura K. MEGA X: molecular evolutionary genetics analysis across computing platforms. Mol Biol Evol. 2018;35:1547–1549.

