## Additional file for "Surfaceome CRISPR Screen Identifies OLFML3 as a Rhinovirus-inducible IFN Antagonist"

### Additional file1: Fig S1-S10

### Table of Content:

Fig. S1. Construction of CRISPR genome-wide and surfaceome libraries.
Fig. S2. Quality analyses of constructed genome-wide and surfaceome CRISPR libraries.
Fig. S3. sgRNA distribution in surfaceome and genome-wide CRISPR libraries post RV-14 challenge.
Fig. S4. T7E1 analyses of the gene modification efficiency of sgRNA for candidate genes identified from surfaceome (A) and genome-wide screens (B).
Fig. S5. RT-qPCR quantification of viral loads in the lysate of knockout cells of the top 10 hits identified from genome-wide screen.
Fig. S6. Representative IF images of knockout cells at 16 h post infection of RV-B14 at an MOI of 1.
Fig. S7. Validation of single clones of ICAM-1$^{-/-}$, RAB5C$^{-/-}$ and OLFML3$^{-/-}$ H1-Hela cells.
Fig. S8. Validation of the effects of ICAM-1, RAB5C and OLFML3 on RV infection, related to Fig. 3.
Fig. S9. Dissection of the functions of RAB5C and OLFML3 in RV infection.
Fig. S10. RNA-Seq analyses of the effects of OLFML3 on RV infection (related to Fig. 4).

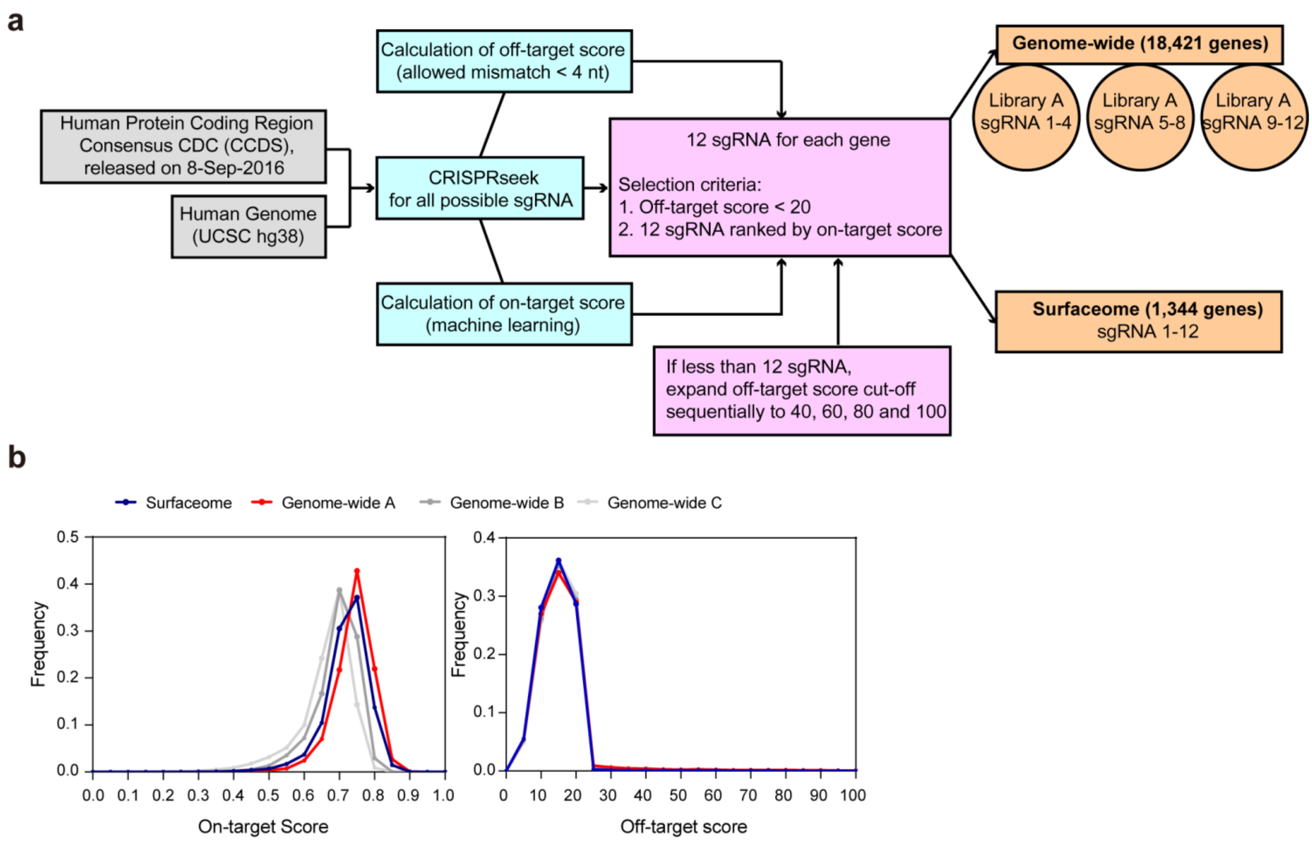


**Fig. S1 Construction of CRISPR genome-wide and surfaceome libraries. a** Schematic illustration. **b** The distribution of sgRNA on-target and off-target scores in genome-wide sub-libraries A, B, C and surfaceome library.

**a**

| | Genome-wide plasmid library | Surfaceome plasmid library | Genome-wide in H1-Hela | Surfaceome in H1-Hela |
|---|---|---|---|---|
| Unique sgRNA in library | 73,527 | 16,975 | 73,527 | 16,975 |
| sgRNA of reads = 0 | 0 | 0 | 0 | 0 |
| Total Reads | 45,880,716 | 8,588,600 | 45,520,228 | 46,194,920 |
| Mapped reads | 38,678,513 | 6,579,411 | 38,796,554 | 32,621,616 |
| Mapping ratio | 84.30% | 76.61% | 85.23% | 70.62% |
| log (Gini index) | 0.042 | 0.029 | 0.051 | 0.034 |

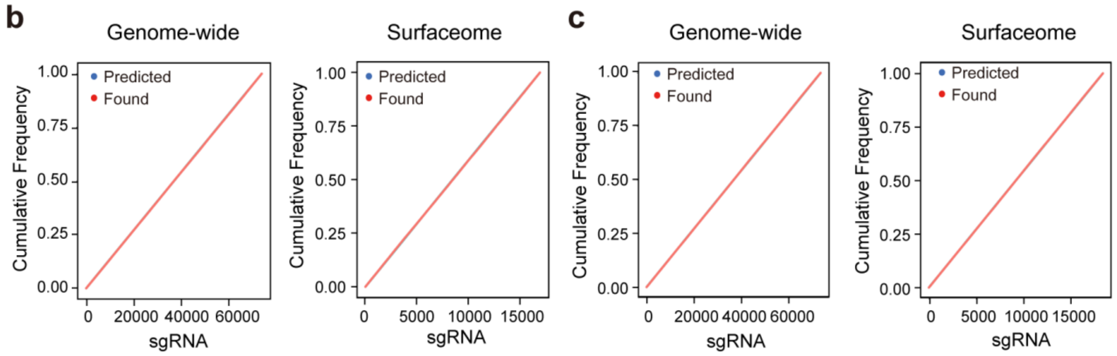


**Fig. S2 Quality analyses of constructed genome-wide and surfaceome CRISPR libraries. a** Summary of next-generation sequencing results of sgRNA in the genome-wide and surfaceome plasmid and H1-Hela libraries. **b**-**c** Distribution of sgRNA in genome-wide and surfaceome libraries in pooled plasmids (**b**) and H1-Hela cells (**c**).

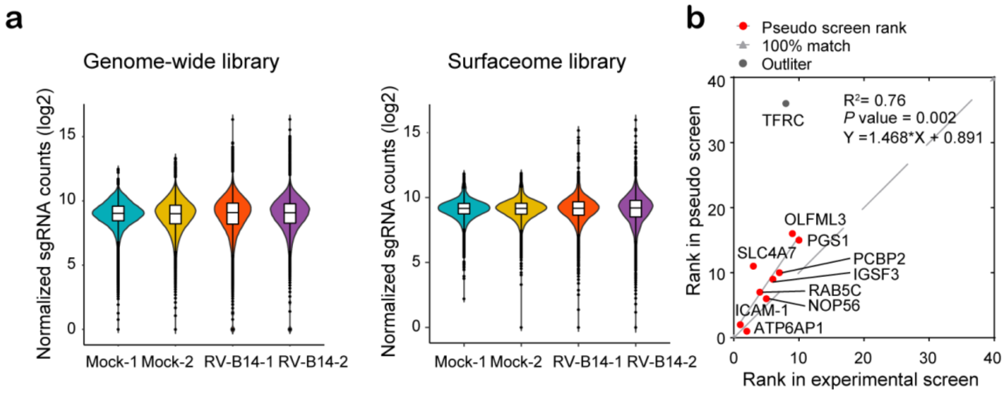


**Fig. S3 sgRNA distribution in surfaceome and genome-wide CRISPR libraries post RV-14 challenge. a** Experimental results of sgRNA enrichment. **b** Correlation analyses of the top 10 hits from experimental surfaceome screen and their ranks in *in silico* (pseudo) surfaceome screen using sgRNA 1-4. The R square and *P* values of the correlation are calculated and shown.

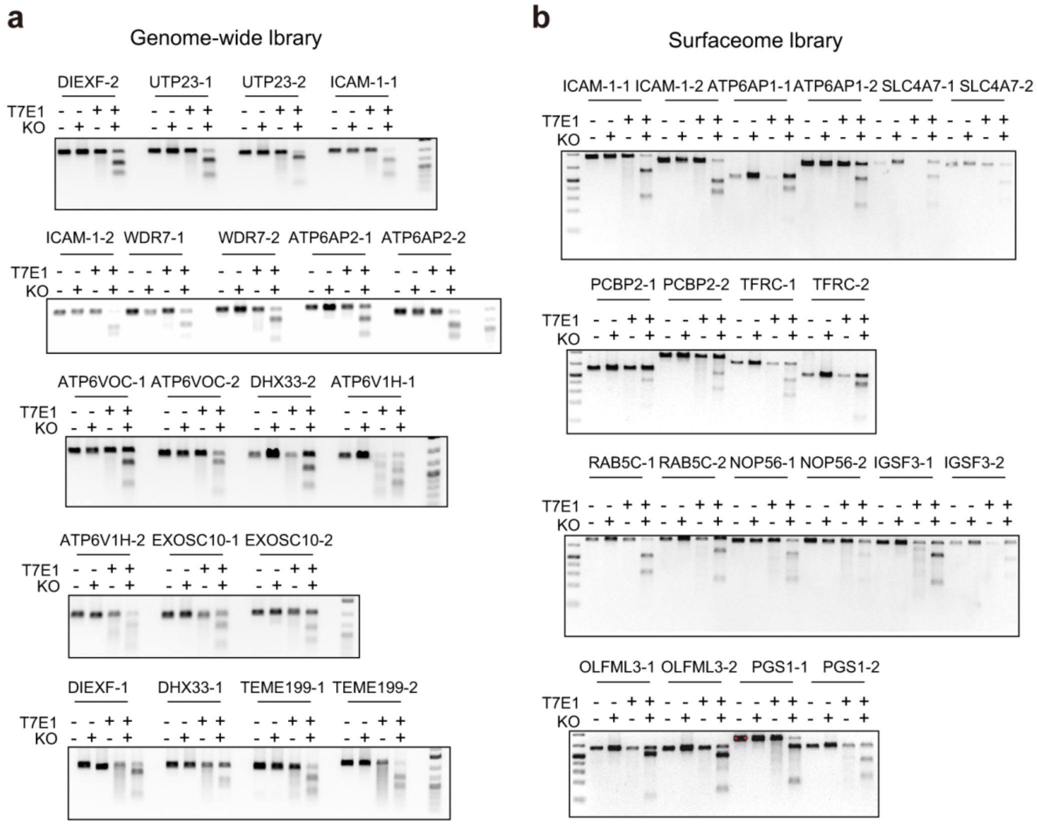


**Fig. S4 T7E1 analyses of the gene modification efficiency of sgRNA for candidate genes identified from surfaceome (a) and genome-wide screens (b).**

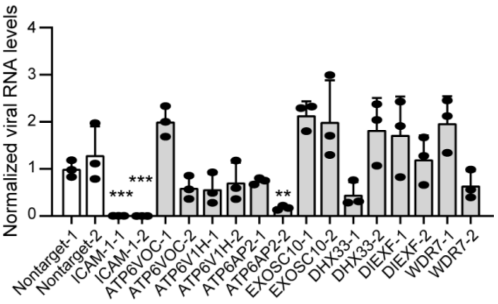


**Fig. S5 RT-qPCR quantification of viral loads in the lysate of knockout cells of the top 10 hits identified from genome-wide screen.** Cell lysate is harvested at 24 h post RV-B14 infection at an MOI of 2. Viral RNA is normalized to RPLP0. The significant difference between mock and knockout cells is determined using two-tailed unpaired Student's *t*-test, $P$ = 0.0006 for ICAM-1-1 and ICAM-1-2, and $P$ = 0.0015 for ATP6AP2-2.

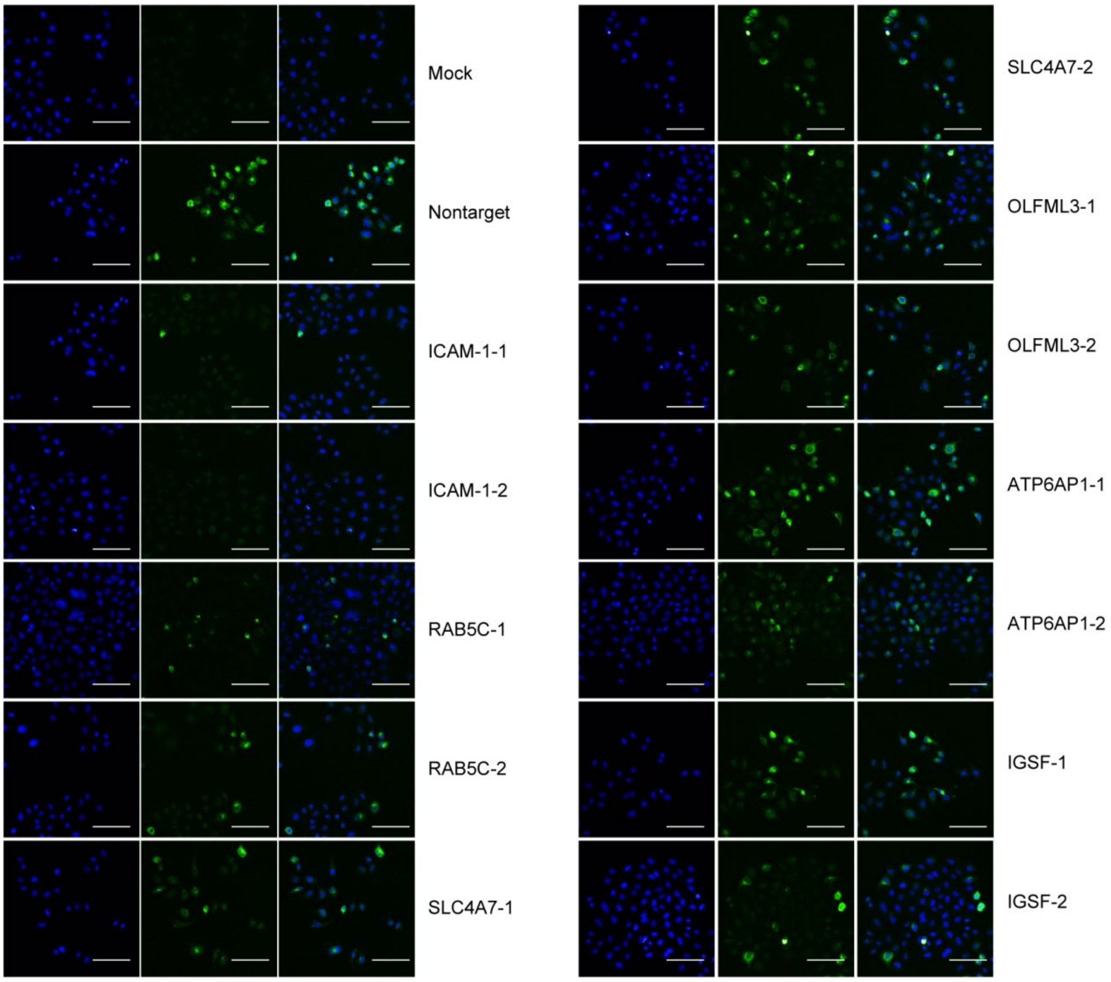


**Fig. S6 Representative IF images of knockout cells at 16 h post infection of RV-B14 at an MOI of 1.**

DAPI, blue; RV-B14 envelope protein, green. Scale bar, 100 μm.

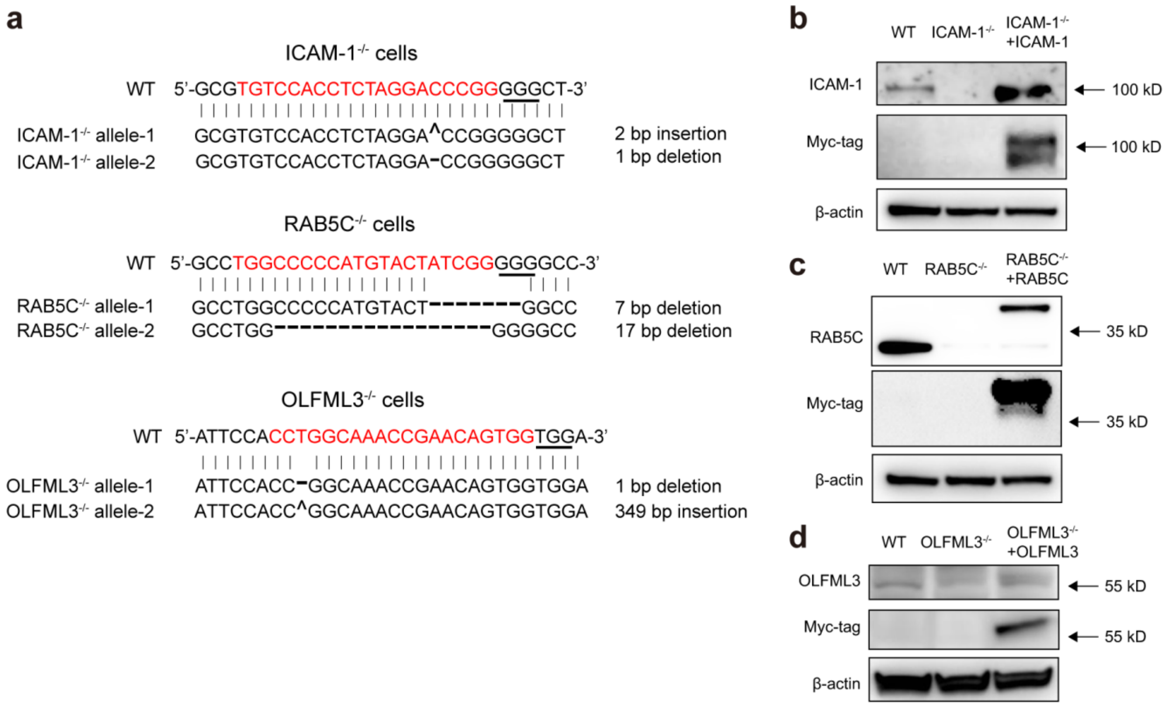


**Fig. S7 Validation of single clones of ICAM-1$^{-/-}$, RAB5C$^{-/-}$ and OLFML3$^{-/-}$ H1-Hela cells. a** Sanger sequencing analyses of mutated alleles. The 20-bp CRISPR-Cas9 target sequences are highlighted in red and protospacer adjacent motif (PAM) underlined. **b-d** Western blot analyses of ICAM-1 (**b**), RAB5C (**c**) and OLFML3 (**d**) expression in wide-type, knockout and rescued cells.

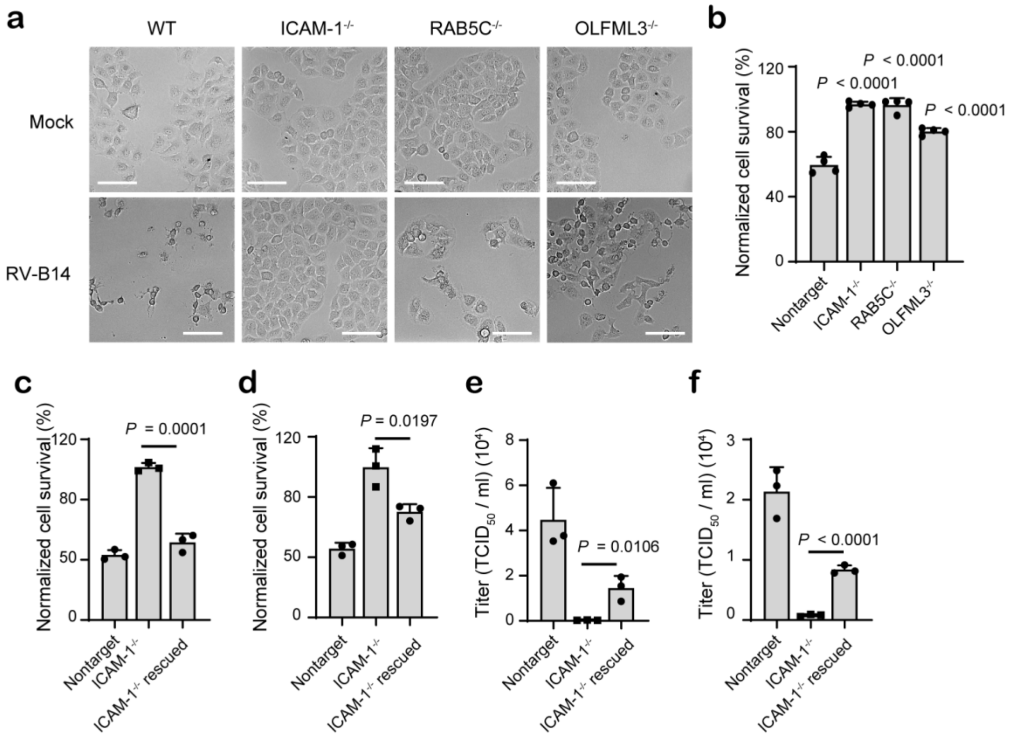


**Fig. S8 Validation of the effects of ICAM-1, RAB5C and OLFML3 on RV infection, related to Fig. 3.**

**a**-**b** Cell viability assay to determine the protective effects of ICAM-1, RAB5C and OLFML3 knockout against RV-B14 infection. Experiments are performed with an MOI of 2 and cell viability is determined at 24 h post infection. **a** Representative images. Scale bar, 100 μm. **b** Quantification of cell viability. Significant difference between test groups and non-targeting sgRNA group is determined using two-tailed Student's *T* test and the *P* values are shown. **c-d** Cell viability assay to determine the effects of ICAM-1 overexpression on RV-B14 (**c**) or RV-A16 (**d**)-induced cell death in ICAM-1$^{-/-}$ cells. **e-f** Rescued susceptibility of ICAM-1$^{-/-}$ H1-Hela cells to RV-B14 (**e**) and RV-A16 (**f**) infection by ICAM-1 overexpression, as determined by viral loads in medium supernatant. Significant difference between knockout and rescued cells is determined using two-tailed unpaired Student's *t* test.

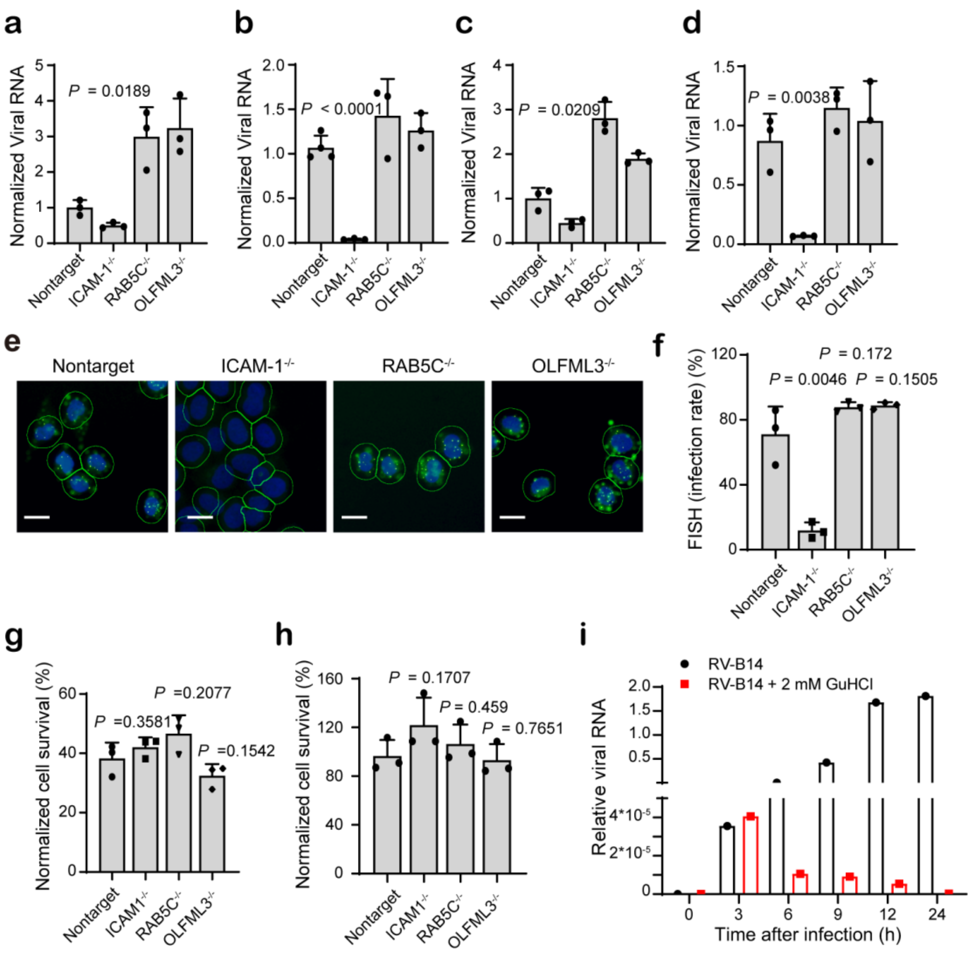


**Fig. S9 Dissection of the functions of RAB5C and OLFML3 in RV infection. a-b** RV-B14 attachment (**a**) and entry (**b**) assays. **c-d** RV-A16 attachment (c) and entry (d) assays. For **a-d**, attached or internalized RV RNA is normalized to RPLP0. **e** Representative images. DAPI, blue; viral genome RNA, green dots; cell membrane, green lines. Scale bar, 20 μm. **f** Quantification of FISH experiments. The results are shown as mean ± SD ($n$ = 3). In each replicate, 5,000 cells are analyzed. **g** Cell viability of mock and knockout cells at 24 h after transfection of RV-A16 genome RNA. **h** Cell viability at 6 h after treatment with 2 mM GuHCl. Significant difference between mock and knockout cells is determined using two-tailed unpaired Student's $t$ test. **i** Inhibition of the synthesis of RV-B14 viral RNA by treatment with 2 mM GuHCl. Cells are infected with RV-B14 at an MOI of 20 and viral RNA in cell lysates is determined and normalized to RPLP0.

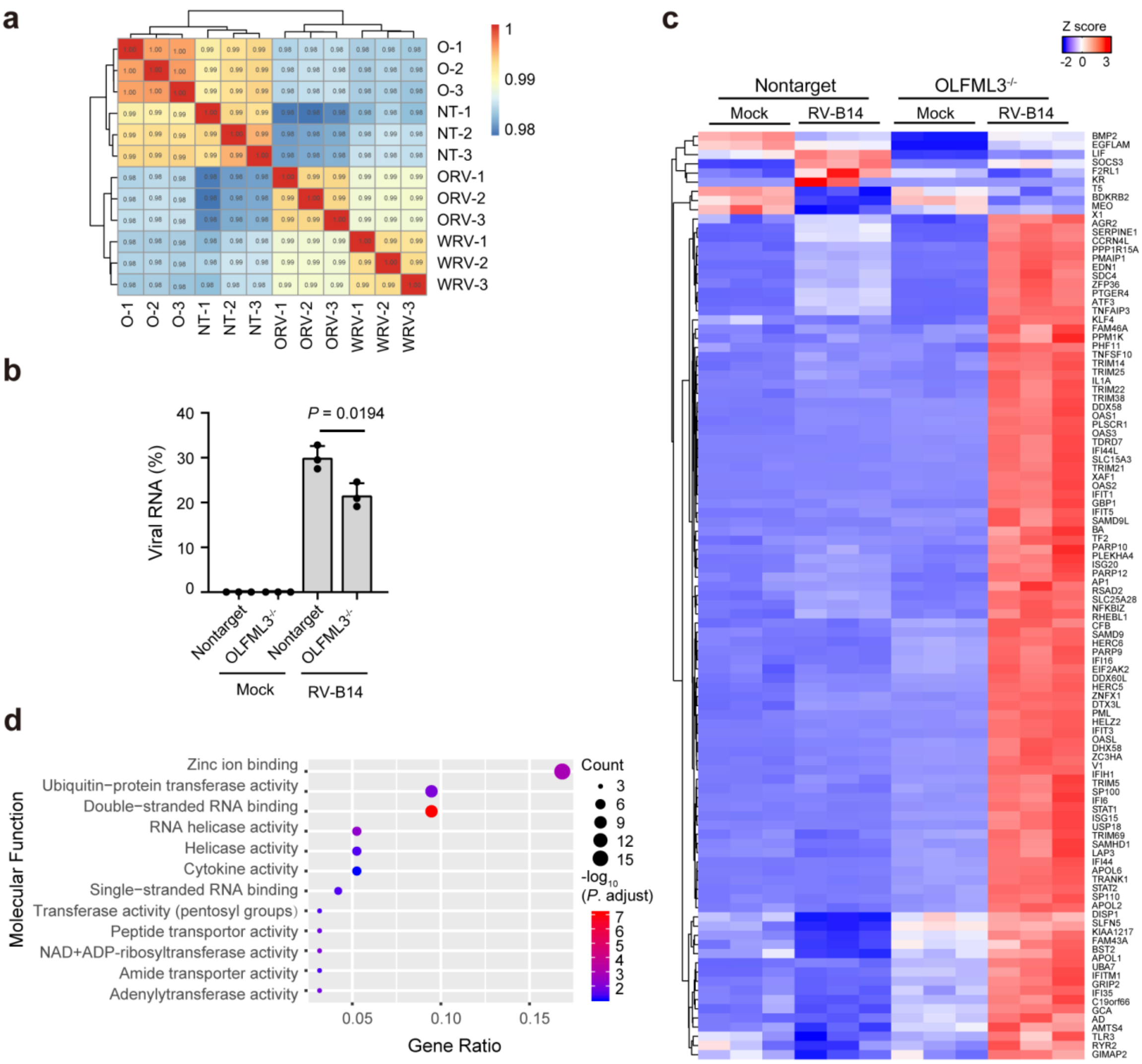


**Fig. S10 RNA-Seq analyses of the effects of OLFML3 on RV infection (related to Fig. 4). a** Pearson correlation analyses of sequenced samples. The symbols are NT for non-targeting sgRNA-transduced cells, O for OLFML3$^{-/-}$ cells and RV for RV-B14 infection. **b** Analyses of the effects of OLFML3 knockout on the transcriptomic expression of RV-B14. The significant difference of RV transcriptomic expression between mock and OLFML3$^{-/-}$ cells is determined using two-tailed unpaired Student's *t* test. **c** Heat map showing the differentially expressed genes with adjusted *P* values of less than 0.05and fold change of more than 2. Cells are collected for RNA-Seq analyses at 24 h after infection with RV-B14 at an MOI of 2. **d** GO analyses of the molecular functions of DEGs.
