## Additional file 2 for "Surfaceome CRISPR Screen Identifies OLFML3 as a Rhinovirus-inducible IFN Antagonist"

### Additional file 1: Table S1-S6

### Table of Content:

Table S1. Primers for construction of CRISPR library and RNA-Seq analyses.
Table S2. Primers for construction of sgRNA plasmid
Table S3. Primers for PCR amplification of sgRNA targeted sites for T7E1 analyses
Table S4. RV-B14-specific or RV universal primers for Taqman RT-qPCR
Table S5. Primers for RT-qPCR
Table S6. siRNA sequences

**Table S1. Primers for construction of CRISPR library and RNA-Seq analyses**

| Primers | Sequences |
|---|---|
| Lib-F | 5’-gtaacttgaaagtatttcgatttcttggctttatatatcttgtggaaaggacgaaacacc-3’ |
| Lib-R | 5’-actttttcaagttgataacggactagccttattttaacttgctatttctagctc-3’ |
| NGS-F | 5’-tctttccctacacgacgctcttccgatctccgtaacttgaaagtatttcga-3’ |
| NGS-R | 5’-gtgactggagttcagacgtgtgctcttccgatctctttttcaagttgataacggac-3’ |

**Table S2. Primers for construction of sgRNA plasmid**

| Genes | Forward primers | Reverse primers |
|---|---|---|
| ICAM1-1 | caccgtgtccacctctaggacccgg | aaacccgggtcctagaggtggacac |
| ICAM1-2 | caccgtgagattgtcatcatcactg | aaaccagtgatgatgacaatctcac |
| ATP6AP1-1 | caccggtgtcattgtaactcacagg | aaaccctgtgagttacaatgacacc |
| ATP6AP1-2 | caccggacctgggaagcattgaagt | aaacacttcaatgcttcccaggtcc |
| SLC4A7-1 | caccgcaccaaggattatctcccag | aaacctgggagataatccttggtgc |
| SLC4A7-2 | caccgccagcatgactgttccattg | aaaccaatggaacagtcatgctggc |
| RAB5C-1 | caccgtggcccccatgtactatcgg | aaacccgatagtacatgggggccac |
| RAB5C-2 | caccgctggcccccatgtactatcg | aaaccgatagtacatgggggccagc |
| NOP56-1 | caccgcgagagcttctccagtcgtg | aaaccacgactggagaagctctcgc |
| NOP56-2 | caccgagccccaaggatctgcactg | aaaccagtgcagatccttggggctc |
| IGSF3-1 | caccgggaatggtaccggctgacgg | aaacccgtcagccggtaccattccc |
| IGSF3-2 | caccgatacggtacttacgccgagg | aaaccctcggcgtaagtaccgtatc |
| PCBP2-1 | caccgtttgtcaatgatcatagcaa | aaacttgctatgatcattgacaaac |
| PCBP2-2 | caccgtggacggcttgggccggtac | aaacgtaccggcccaagccgtccac |
| TFRC-1 | caccgttgttagtctggaagtagca | aaactgctacttccagactaacaac |
| TFRC-2 | caccgcggagccccagaagacatgt | aaacacatgtcttctggggctccgc |
| OLFML3-1 | caccgccactgttcggtttgccagg | aaaccctggcaaaccgaacagtggc |
| OLFML3-2 | caccgttgggctgtctatgccaccc | aaacgggtggcatagacagcccaac |
| PGS1-1 | caccggatcctgctggcctcaccag | aaacctggtgaggccagcaggatcc |
| PGS1-2 | caccgtgatggcatccctctacctg | aaaccaggtagagggatgccatcac |
| DIEXF-1 | caccgcatgcaggcaatacacatgg | aaacccatgtgtattgcctgcatgc |
| DIEXF-2 | caccgactacccctggactcacatg | aaaccatgtgagtccaggggtagtc |
| UTP-23-1 | caccggctgccccgctacctcatgg | aaacccatgaggtagcggggcagcc |
| UTP-23-2 | caccgacagctgcgtctcccccatg | aaaccatgggggagacgcagctgtc |
| WDR7-1 | caccgtgccatcactcacatcccag | aaacctgggatgtgagtgatggcac |
| WDR7-2 | caccgagccgcgcagactatcacca | aaactggtgatagtctgcgcggctc |
| ATP6AP2-1 | caccgaagtaccatgttgaaaacca | aaactggttttcaacatggtacttc |
| ATP6AP2-2 | caccgaggagagcggatcccagacg | aaaccgtctgggatccgctctcctc |
| ATP6V0C-1 | caccggggcgacgatgagaccgtag | aaacctacggtctcatcgtcgcccc |
| ATP6V0C-2 | caccgcaagagcggtaccggcattg | aaaccaatgccggtaccgctcttgc |
| DHX33-1 | caccgctgctaccggctctacacgg | aaacccgtgtagagccggtagcagc |
| DHX33-2 | caccgtctgcgtcatagatgcacag | aaacctgtgcatctatgacgcagac |
| ATP6V1H-1 | caccggatgtagcaagaacactgcg | aaaccgcagtgttcttgctacatcc |
| ATP6V1H-2 | caccgacgatgttggagaatatgtg | aaaccacatattctccaacatcgtc |
| TMEM199-2 | caccgctcggccagacccaccacag | aaacctgtggtgggtctggccgagc |

| | | |
|---|---|---|
| TMEM199-3 | caccgtccaaggtaagtgcagacga | aaactcgtctgcacttaccttggac |
| EXOSC10-1 | caccgcagcaagctatgatgccctg | aaaccagggcatcatagcttgctgc |
| EXOSC10-2 | caccgcctgaaactctactgcaacg | aaaccgttgcagtagagtttcaggc |
| ITPK1-1 | caccgtgagctcataggacttggag | aaacctccaagtcctatgagctcac |
| ITPK1-2 | caccgtcatcatcaacaaccagaca | aaactgtctggttgttgatgatgac |

**Table S3. Primers for PCR amplification of sgRNA targeted sites for T7E1 analyses**

| Genes | Forward primers | Reverse primers |
|---|---|---|
| ICAM1-1 | ggagaaggagctgaaacggg | gtgtctcctggctctggttc |
| ICAM1-2 | gctgttcccagtctcggagg | ggggccatctggaaaaacac |
| ATP6AP1-1 | aaggctggttgggtgtttct | acctcctacatcctcacgct |
| ATP6AP1-2 | ggtggttgaggtatgggtgg | cacacagacacgggtcagaa |
| SLC4A7-1 | agggataggaagaattactgtggc | gttcccctggagcagaacat |
| SLC4A7-2 | acgtgtttggtcgtggttttc | tccttggttcccatgtgttca |
| RAB5C-1 | tcctggcttgagggttctct | gcgagtgcaatgacgatgtt |
| RAB5C-2 | tgtgcttggagtaggcttcc | gatgcaatgtccccaaacagg |
| NOP56-1 | cctccaggcagagtaggtgt | aagacatggtgggtcacgaaa |
| NOP56-2 | tcgtgacccaccatgtcttc | ttaccccttgtcttcagggc |
| IGSF3-1 | ggcccagatgctgtctttgg | ttcccttgtgagtctgaaacgc |
| IGSF3-2 | ggtgattcctgtgttcccct | caggacccactgcaggtaaa |
| PCBP2-1 | cccctttttgggaagtgtggt | gcgagtctcaaggtccagtg |
| PCBP2-2 | catggacatctcccttgatgaga | agaccaccatccagaatctcc |
| TFRC-1 | ccttgaaccaaaatgttttcccc | cagcagaaacagaaaatgacagc |
| TFRC-2 | aaagtcacagggctgtgtaaa | ccctgtattaaaagctgctgcc |
| OLFML3-1 | gttcctgctggaggcatctaa | gtgtctgctgtcaagccgta |
| OLFML3-2 | cagtattcccagcagagggg | agctcctcaaacctcctcct |
| PGS1-1 | gcccctgacacctggattta | cttgaggagatgaatgctcgc |
| PGS1-2 | cttaatggccgtgttgcatt | tggggcaacaaaaacaagagc |
| DIEXF-1 | cagccagttcctatctggtcc | ggctcattaaggtcccatcca |
| DIEXF-2 | ccaggagggccagttttagc | ggagtgggttacctggcatc |
| UTP23-1 | tgcgtgaggcgtttactgat | acgggtacactttggataccg |
| UTP23-2 | ttcttccgcaacaacttcgg | actgtgaggcttgcagagttt |
| WDR7-1 | gaatgtggactatggtggggg | aggacctagagcattttccca |
| WDR7-2 | ctttcctcccaactcccaagt | agtttccatcctttccgtcgaa |
| ATP6AP2-1 | accatcagtgcaagtgtcgt | tccataacacgagctattctaaatg |
| ATP6AP2-2 | aggactcagcattttgaccagt | cagaagccaagcttgaagcac |
| ATP6V0C-1 | atgtcagtcctctcttctcgc | tccatccaggaagtctcagc |
| ATP6V0C-2 | cccaagacctttggtggctt | gctgatgtcgtcattcaggga |
| DHX33-1 | ctgcggagcagtctggtttt | tccatttgttgttgtttgtctcc |
| DHX33-2 | ttggtgaccacctccgaatg | agatgctttcacgctcctgg |
| ATP6V1H-1 | tctgtgcctactaccatccct | tcctatgggaacggacaggtt |
| ATP6V1H-2 | tgcttagtgctcagtccacc | gcaaccggcctatactactca |
| TMEM199-1 | gacagctaaagtcacctgcg | cgggcaactagttctgggtt |
| TMEM199-2 | agagctttccagcagtcacg | gggaaggcctggattccatt |
| EXOSC10-1 | gcttggtgtcgcaaagaaca | acaggcttgctaagttggga |
| EXOSC10-2 | atgggacagcggagactacc | tttcccacctatggaaggtca |
| ITPK1-1 | ctccaggtgactgctcgaaa | cctgccactgagctgtctac |
| ITPK1-2 | cagtaggtctgtctggcttcc | acatgtaaactgcaggggggc |

**Table S4. RV-B14-specific or RV universal primers for Taqman RT-qPCR**

| Genes | Sequence |
|---|---|
| RV-B14-F | ctagcctgcgtgg |
| RV-B14-R | aaacacggacacccaaagt |
| RV-B14-probe | FAM-tcctccggcccctga-MGB-NFQ |
| RV-general-F | gtgaagagccscrtgtgct |
| RV-general-R | gctscagggttaaggttagcc |
| RV-general-probe | fam-tgagtcctccggcccctgaatg-BHQ1 |

Note: Degenerate codons are R for A or G and S for C or G.

### Table S5. Primers for RT-qPCR

| Genes | Forward primers | Reverse primers |
|---|---|---|
| IFIT1 | caatactgggtacgcaatcacc | agctcgttttaggacgtgcag |
| IFIT2 | gacacggttaaagtgtggagg | tccagacggtagcttgctatt |
| IFIT3 | aaaagcccaacaacccagaat | cgtattggttatcaggactcagc |
| OAS1 | tgtccaaggtggtaaagggtg | ccggcgatttaactgatcctg |
| OAS2 | ctcagaagctgggttggtttat | accatctcgtcgatcagtgtc |
| OAS3 | gatgtgctgccagcctttga | tgtagctctgtgaagcaggtgga |
| ISG15 | cgcagatcacccagaagatt | gcccttgttattcctcacca |
| ISG20 | ctcgttgcagcctcgtgaa | cgggttctgtaatcggtgatctc |
| IFNB | gagctacaacttgcttggattc | caagcctcccattcaattgc |
| STAT1 | cagcttgactcaaaattcctgga | tgaagattacgcttgcttttcct |
| STAT2 | ccagctttactcgcacagc | agccttggaatcatcactccc |
| SOCS3 | cctgcgcctcaagaccttc | gtcactgcgctccagtagaa |
| OLFML3 | cgccgactagctgctttagag | cgctccagacgatccactc |

**Table S6. siRNA sequences**

| Genes | Sense sequences (5'-3') | Anti-sense sequences (5'-3') |
|---|---|---|
| SOCS3-1 | accuuucugauccgcgacatt | ugucgcggaucagaaaggutt |
| SOCS3-2 | gccuauuacaucuacuccgtt | cggaguagauguaauaggctt |
